# Genomic signatures of a major adaptive event in the pathogenic fungus *Melampsora larici-populina*

**DOI:** 10.1101/2021.04.09.439223

**Authors:** Antoine Persoons, Agathe Maupetit, Clémentine Louet, Axelle Andrieux, Anna Lipzen, Kerrie W. Barry, Hyunsoo Na, Catherine Adam, Igor V. Grigoriev, Vincent Segura, Sébastien Duplessis, Pascal Frey, Fabien Halkett, Stéphane De Mita

## Abstract

**Background:** The recent availability of genome-wide sequencing techniques has allowed systematic screening for molecular signatures of adaptation, including in non-model organisms. Host-pathogen interactions constitute good models due to the strong selective pressures that they entail. We focused on an adaptive event which affected the poplar rust fungus *Melampsora larici-populina* when it overcame a resistance gene borne by its host, cultivated poplar. Based on 76 virulent and avirulent isolates framing narrowly the estimated date of the adaptive event, we examined the molecular signatures of selection.

**Results:** Using an array of genome scan methods, we detected a single locus exhibiting a consistent pattern suggestive of a selective sweep in virulent individuals (excess of differentiation between virulent and avirulent samples, linkage disequilibrium, genotype-phenotype statistical association and long-range haplotypes). Our study pinpoints a single gene and further a single amino acid replacement which may have allowed the adaptive event. Although the selective sweep occurred only four years earlier, it does not seem to have affected genome diversity further than the immediate vicinity of the causal locus.

**Conclusions:** Our results suggest that *M. larici-populina* underwent a soft selective sweep and possibly a prominent effect of outbreeding and recombination, which we speculate have increased the efficiency of selection.

## Background

Understanding adaptation in response to natural or anthropic forces is one of the major goals of the study of molecular evolution. In most cases, evolutionary research focuses on historical events that occurred in a more or less distant past, such as domestication [1, 2, 3], divergence of populations [4, 5, 6], adaptation to environmental conditions [7, 8, 9] or speciation [10]. In comparison, few biological models allow to address the evolutionary process in contemporary or at least very recent events (but see e.g. [11]). Due to their highly dynamic nature, host-pathogen interactions offer opportunities to study adaptation processes at work. Indeed, host-pathogen interactions evolve constantly and rapidly, due to strong and reciprocal selection pressures, changing environmental conditions and, recently, anthropic interactions [12, 13]. Microorganisms pathogenic to plants are considered as capable of extremely fast evolution [14, 15, 16].

It has been shown that the fate of an infection depends on a complex interplay of mechanisms involving both partners of the interaction. The pathogen produces molecules manipulating the host to allow entry and co-optation of resources, but also aiming at preventing the activation of the host defence machinery [17, 18]. Plants have developed both non-specific and specific immunity systems to detect pathogens and prevent infection [19]. Plant immunity systems are based on resistance genes which target molecules characteristic of a given pathogen species or even strain. Molecules produced by pathogens and enabling host defence responses (thereby preventing successful infection) are named avirulence factors. In contrast with non-specific immunity, such host resistance genes allow complete resistance against the targeted pathogens. Selective pressures on genes controlling these systems can be strong and lead to fast co-evolution [20, 21]. In particular, genes coding for proteins interacting directly to determine complete immunity, such as pairs constituted of resistance proteins and avirulence factors, are likely to be exposed to the strongest selective pressures [19, 22].

The strength of selective pressure caused by total host resistance can be extremely high if resistance entails a substantial reduction of ecological niche. This is typically the case when disease-resistant hosts are dominant in the environment. Then a mutation restoring virulence is expected to undergo an extremely fast increase in frequency toward fixation, resulting in a strong selective sweep of genomic variation. However, experimental evidence and modelling suggest that relatively mild selective sweeps could be expected [23], because selection of variants already segregating in the population (soft selective sweep) is expected to be actually faster than selection of a newly occurring mutation (hard selective sweep).

Identifying genomic regions subjected to selection has long been of interest to evolutionary biologists [24]. The recent emergence of highthroughput sequencing technologies offering a genome-wide coverage [25] provides opportunities to detect loci involved in adaptation. Modern sequencing techniques allow to dramatically increase both the number of loci characterized and the number of individuals used in population samples. The rationale underlying the search for the signatures of adaptation in molecular data has been proposed long before genome sequencing became routine [26], while more recent methodologies have been introduced to leverage the wealth of data that became available thanks to whole-genome sequencing. Essentially it boils down to scanning the genome to pinpoint genes or regions exhibiting characteristic footprints left by selective sweeps [27, 28]. These footprints include a deformation of the allele frequency spectrum, an excess of linkage disequilibrium and a reduction in genetic diversity around the selected region. It can also include an excess of differentiation in allelic frequencies between individuals from different subpopulations (e.g. if the sweep only affected a single subpopulation). Note that any one of these can be potentially mimicked by demographic effects such as non-equilibrium or undocumented substructure that can result in an inflated rate of false or true positives, especially for methods that rely on an assumed model [29]. We can expect however that combining different complementary methods looking for different genomic signatures can increase power, reduce sensitivity to confounding factors (which are unlikely to affect different methods in a similar manner) and increase precision of the detection of selective sweeps [30]. Finally if one can show a statistical association between a selected trait and a genotype (like in a QTL or a GWAS analysis), this provides powerful complementary evidence.

We propose to adopt this approach to detect the determinants of a recent adaptation event in a plant pathogenic fungus. We focus on the poplar rust fungus, *Melampsora larici-populina*, which is a major threat for cultivated poplar in Europe [31]. *Melampsora larici-populina* needs two hosts from unrelated genera to complete its life cycle: *Populus* on which it performs several asexual reproduction cycles during summer and autumn and *Larix* on which it performs a single sexual reproduction cycle once a year in spring. *Melampsora larici-populina* is particularly damaging on cultivated poplar hybrids, mostly because of their intensive monoclonal cultivation over several decades [32]. Many poplar cultivars carrying qualitative resistances were bred, but *M. larici-populina* eventually overcame all resistances. The most significant resistance break-down event occurred in 1994 and targeted the RMlp7 resistance, which was at the time carried by most cultivated poplars in Northern France and Belgium. This event has led to the invasion of France by virulent individuals in less than five years [33].

In this article we focus on this resistance and we denote *M. larici-populina* isolates that can successfully infect poplar hosts harbouring RMlp7 as virulent, as opposed to avirulent isolates.

An analysis of the French population structure of *M. larici-populina* showed a strong impact of the resistance breakdown [34]. The analysis of 594 isolates from a laboratory collection sampled before and after the resistance breakdown identified three genetic groups. These groups have been named ‘wild’ (isolates sampled in a region where cultivated poplars are sparse), ‘fossil’ (isolates sampled before the breakdown and shortly afterwards) and ‘cultivated’ (isolates sampled only after the breakdown). All isolates from the two former groups were found to be avirulent and nearly all isolates from the latter group were found to be virulent. Furthermore, signatures of population expansion have been detected in the ‘cultivated’ group over time [34].

In this study we selected 76 isolates from the three genetic groups found in [34], spread into four samples corresponding each to a given year. We defined two samples in the ‘cultivated’ (virulent) group, one immediately after the resistance breakdown and another four years after, after completion of the putative selective sweep (fixation of the virulence allele in the *M. larici-populina* population). Using over a million single-nucleotide polymorphisms (SNPs), we conducted several genome scan approaches to detect specific regions of the genome potentially involved in virulence. Genome scan methods consistently highlighted a common candidate, which is the first ever described putative avirulence gene in *M. larici-populina*. However, in spite of the potential strength of selective pressure to overcome resistance, molecular diversity indicates a relatively mild population bottleneck and the signature of selective sweep are rather locally restricted.

## Results

### Sampling and polymorphism detection

In order to characterize the resistance break-down, 76 isolates were selected from a previous population genetics study investigating temporal evolution of *Melampsora larici-populina* strains throughout France (Additional file 1: Table S1). Four samples framing the breakdown event were defined based on the population structure described in [34]. Each sample was constituted of isolates sampled the same year, though not necessarily from the same location: a 1993 sample (n=18) from the ‘fossil’ group, a 1994 sample (n=18) from the ‘cultivated’ group, a 1998 sample (n=21) from the ‘cultivated’ group and a 2008 sample (n=19) from the ‘wild’ group. Note that *M. larici-populina* isolates are isolated at a stage of their life cycle where they are dikaryotic (two unmerged nuclei per spore cell) and can therefore be treated as diploid. The virulence profile of those isolates was determined and showed that all isolates from the 1993 and 2008 samples were avirulent while all isolates from the 1994 sample were virulent as well as all but one from the 1998 sample (Additional file 1: Table S1).

The genome of those 76 *M. larici-populina* isolates was sequenced with the Illumina technology (150 bp paired reads). An average of 68 million reads were generated per isolate (ranging from 26 to 143 million reads). On average, 78% of the reads aligned on the chromosomes of version 2 of the reference genome [35] generating a sequencing depth of 80X (± 24) (Additional file 1). In total, the site calling procedure identified 81,174,498 fixed sites and 1,125,506 SNPs covering more than 80% of the total length of the 18 chromosomes (Additional file 2: Table S2). The remaining 19% of sites were not called due to missing data or, much less frequently, sites with more than two alleles. The amount of variation is higher than previously identified [36], with more than 1% of sites with enough data exhibiting polymorphism.

### Genetic structure

To assess the genetic structure of our dataset, we performed a non-parametric structure analysis based on a discriminant analysis of principal components (DAPC) [37]. The number of genetic groups was assessed based on the Bayesian information criterion (BIC) values (Additional file 3: Fig. S1a). The BIC criterion points to two groups, gathering on one hand the 1993 and 2008 samples (avirulent cluster) and on the other hand the 1994 and 1998 samples (virulent cluster; Additional file 3: Fig. S1b). We analysed the results with three and four groups as well, due to the proximity of BIC values. With three groups, the avirulent cluster is split in two, separating the 1993 and 2008 groups (Additional file 3: Fig. S1c), in a structure identical to [34]. With four groups, a part of 1994 isolates are separated from the virulent cluster, keeping the other 1994 isolates together with the whole 1998 sample (Additional file 3: Fig. S1d). This group of isolates seems to be only partially related to the region of origin of samples since it includes all 5 isolates sampled in Belgium, 2 of the 8 isolates sampled in Nord-Pas-de-Calais, 1 of the 3 isolates sampled in Picardie and the single isolate sampled in Centre. Up to four groups, all assignation coefficients in DAPC analyses are virtually equal to 1 (Additional file 3: Fig. S1e-g).

In addition, we reconstructed a neighbour-joining tree based on pairwise distances computed from all SNPs (Fig. 1). This tree distinguishes three clades: the 2008 sample, the 1993 sample and a third clade gathering 1994 and 1998 samples. These clades exhibit a star-like topology, especially the 2008 and 1993 samples (with the exception of a pair of 1993 samples which exhibit high pairwise similarity). The 1994 and 1998 samples are interspersed within the third clade that exhibits comparatively longer internal branches. The group of 1994 samples that forms a group of its own in the DAPC analysis with four groups forms a subclade, although it does not appear to show marked pairwise divergence with respect to the other isolates.

**Figure 1:**
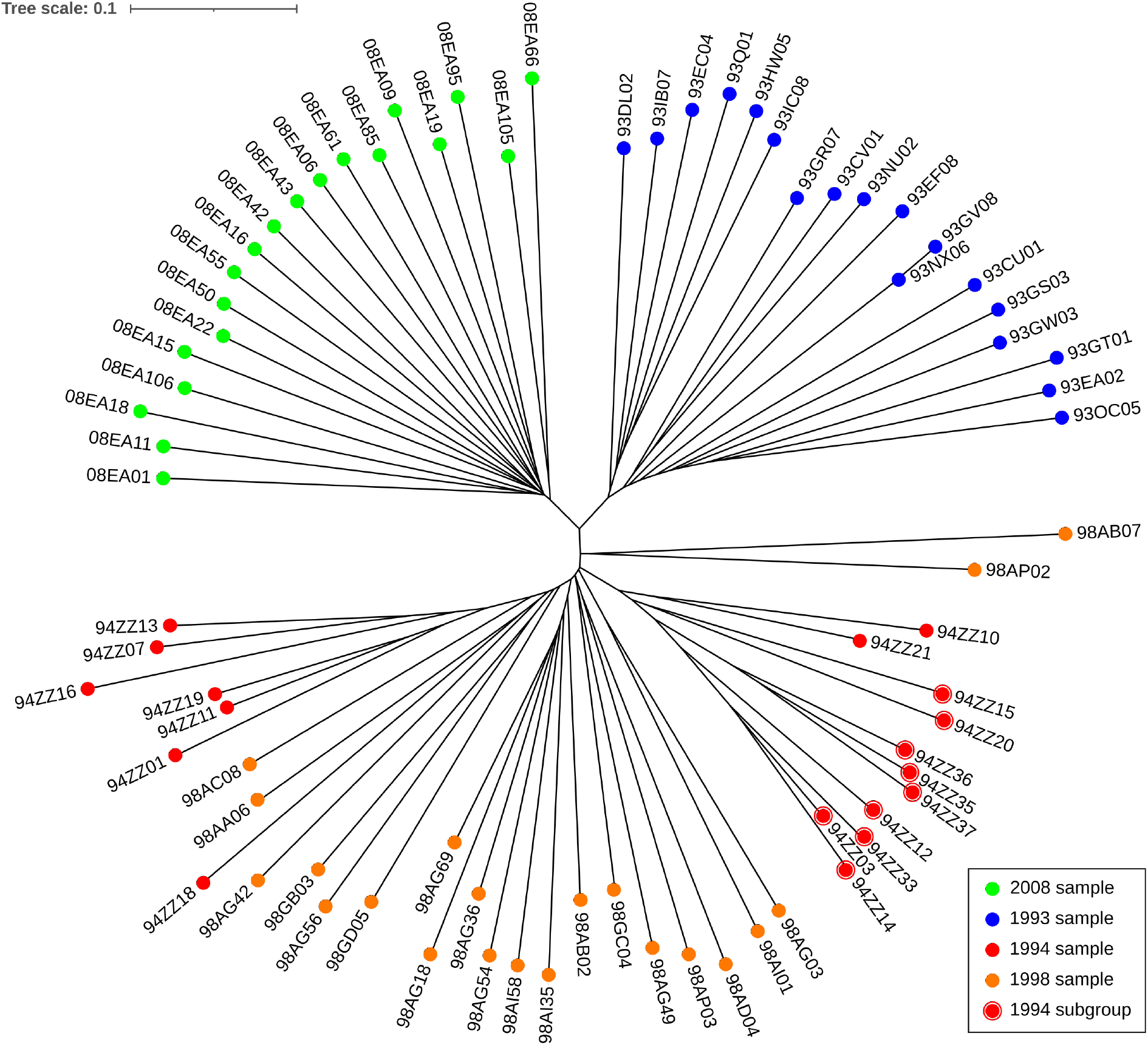
Unrooted neighbour-joining tree of all isolates based on genomic distances. The distances are expressed in number of differences per compared position (due to missing data, the number of compared positions is less than the number of polymorphic sites and varies between comparisons). Coloured disks indicate sample membership. Isolates of the 1994 sample that are assigned to a specific group by the DAPC analysis are indicated by an additional red circle.

The analysis of genome-wide statistics measuring pairwise differentiation between samples gives results consistent with the tree structure (Additional file 4: Table S4i-n). The virulent cluster (1994 and 1998) is grouped with strong similarity (*D*_*a*_ = 0.0035), as well as the avirulent cluster (1993 and 2008), but with a lesser similarity (*D*_*a*_ = 0.0109). Between these two clusters, the pairwise comparison results might have been influenced with the presence of substructure within some of the samples with in particular less similarity between samples 1993 and 1994 than between 1993 and 1998 (*D*_*a*_ = 0.0155 and 0.0142, respectively).

The separation of the virulent and avirulent clusters therefore seems to be the main feature of population structure, but it actually represents a small proportion of the total genetic variance. Weir and Cockerham’s estimation of the between-cluster fixation index 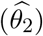 gives a value of 4% of the total variance (Additional file 4: Table S4a). In addition, population substructuration is visible in all but one (2008) sample and especially strong for 1994.

### Linkage disequilibrium

Next we investigated patterns of linkage disequilibrium (LD). Since we used unphased data, all LD values are based on genotypes (individual level instead of haplotype). LD is influenced by recombination rate (which can vary substantially across genomes, but which we don’t expect to vary significantly between samples) and by demographic or selective forces. In our case, we expected an effect of the adaptive event which is expected to increase LD for virulent isolates due to the population bottleneck and fast population growth rate. We compared the LD decay with physical distance between the four samples (Additional file 5: Fig. S2). We found that the average LD drops below *r*^2^ = 0.1 before 95-165 Kbp according to the sample. The 2008 sample appears to be the one where LD drops at the fastest rate and 1994 at the slowest rate (average *r*^2^ below 0.2 before 33 kb for 2008 and before 60 kb for 1994; average *r*^2^ of 0.0955 at 100 kb for 2008 and 0.1459 for 1994), the 1998 and 1993 samples having a slightly slower rate of decay than 2008 (Additional file 5: Table S5). These observations are compatible with the hypothesis that the population from which the 2008 sample has been collected is at equilibrium. In contrast, the virulent population may have undergone an expansion following the resistance breakdown, with the 1994 isolates displaying signatures of a recent adaptive event, which is attenuated in 1998. Potential substructure in the 1994 and 1998 samples may also explain long-distance linkage disequilibrium.

A survey of the level of pairwise linkage at the chromosome level was performed (Additional file 6: Fig. S3). The analysis highlights numerous blocks with extended LD that are shared by several (and often all) samples, such as, for chr01, a region between 5.5 and 6.0 Mbp. At a genome-wide scale, it is visible that the overall levels of LD are more elevated in sample 1994 and lower in sample 2008.

### Population genomics statistics

At a genome-wide scale, statistics describing nucleotide polymorphism capture the consequences of demographic history. Besides the population similarity statistics addressed in a previous section, we computed statistics characterizing the levels and structure of polymorphism (Table 1; see also Additional file 4: Table S4 for the full list). We found substantial levels of diversity, with both 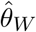 and *π* around 0.002 per site, or the whole dataset as well as for the four samples. The 2008 sample shows the highest value of 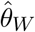 which reflects a larger number of polymorphic sites, but not a high nucleotide diversity (*π*) compared with other samples, showing a marked unbalance of allelic frequencies (with an excess of singletons and low-frequency variants), as illustrated by the strongly negative values for the three considered neutrality tests Tajima’s *D* and Fu and Li’s *D*^∗^ and *F*^∗^, as well as by the marked star-like structure of the tree (Fig. 1). This excess of rare alleles might be explained by recent population expansion, e.g. following a population bottleneck. The expansion might still be older than the bottleneck that affected virulent isolates, since negative values of neutrality tests indicate that polymorphism has started to regenerate.

**Table 1:**
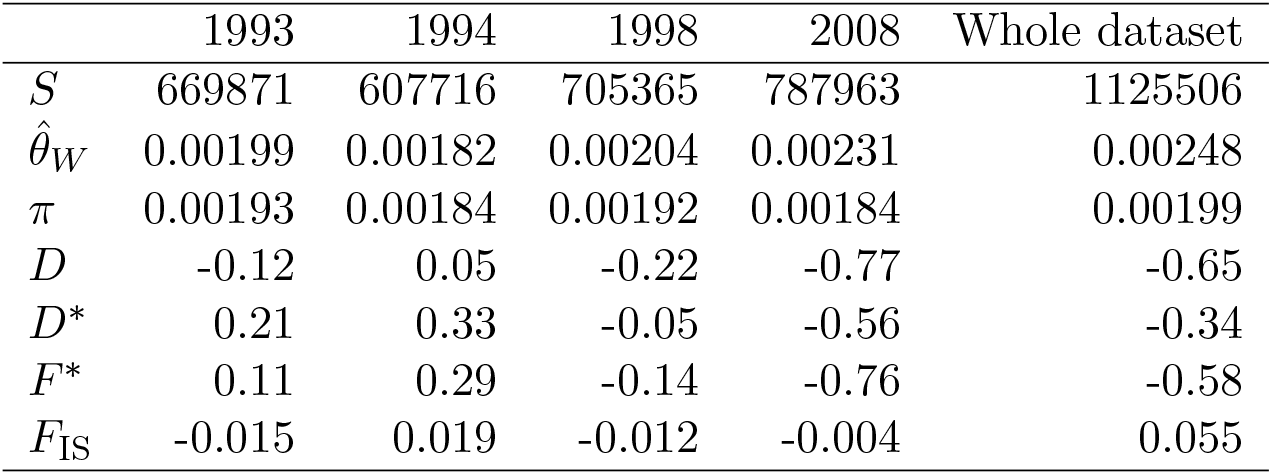
Genome-wide diversity statistics. Statistics are defined in Additional file 4: Table S3

We expected to find signatures of a strong on-going bottleneck in sample 1994 and of population expansion in 1998. Features of polymorphism are compatible with these expectations but they point to a weak or moderate intensity. Compared with the 1993 sample, the 1994 sample exhibits reduced levels of diversity with a more marked reduction for 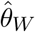 (about 10%) than *π* (about 5%) and, consequently, positive values for Tajima’s and Fu and Li’s tests of neutrality. This is the expected pattern during a bottleneck. Between 1994 and 1998, diversity increased consistently. The comparatively large increase of 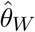 is reflected by the negative values of neutrality tests and points to a relatively high mutation rate.

*F*_IS_ values indicate a substantial deficit of heterozygotes in the whole dataset, likely due to sample structure. With the exception of the 2008 sample, we note deviations in both directions, ranging from −1.2 to +1.9%, with respect to Hardy-Weinberg equilibrium. This is yet another signature indicating that the populations we have sampled were likely not at the equilibrium and/or exhibit potential substructure.

### Localisation of a avirulence gene candidate

In order to identify the locus or loci involved in the resistance breakdown, we compared the results of several selective sweep detection methods addressing different signatures of a selective sweep event. These signatures fall into three main categories: deformation of the allele frequency spectrum, excess of differentiation between virulent and avirulent isolates and extended linkage disequilibrium.

### Allele frequency spectrum

The effect of selection on the allele frequency spectrum should be captured by Tajima’s *D*. We computed Tajima’s *D* on a sliding window separately for avirulent and virulent isolates (Fig. 2, first two panels) as well as for all samples (Additional file 7: Fig. S4) but did not find any clear peak anywhere in the genome. A Bayesian, hidden Markov model method aiming to detect selective sweeps (freq-hmm) was also applied on the 1998 sample (when the selective sweep should be completed). We detected 22 potential selective sweeps on 11 out of 18 chromosomes, containing together a total of 614,822 SNPs (Fig. 2, third panel, Additional file 8: Table S5).

**Figure 2:**
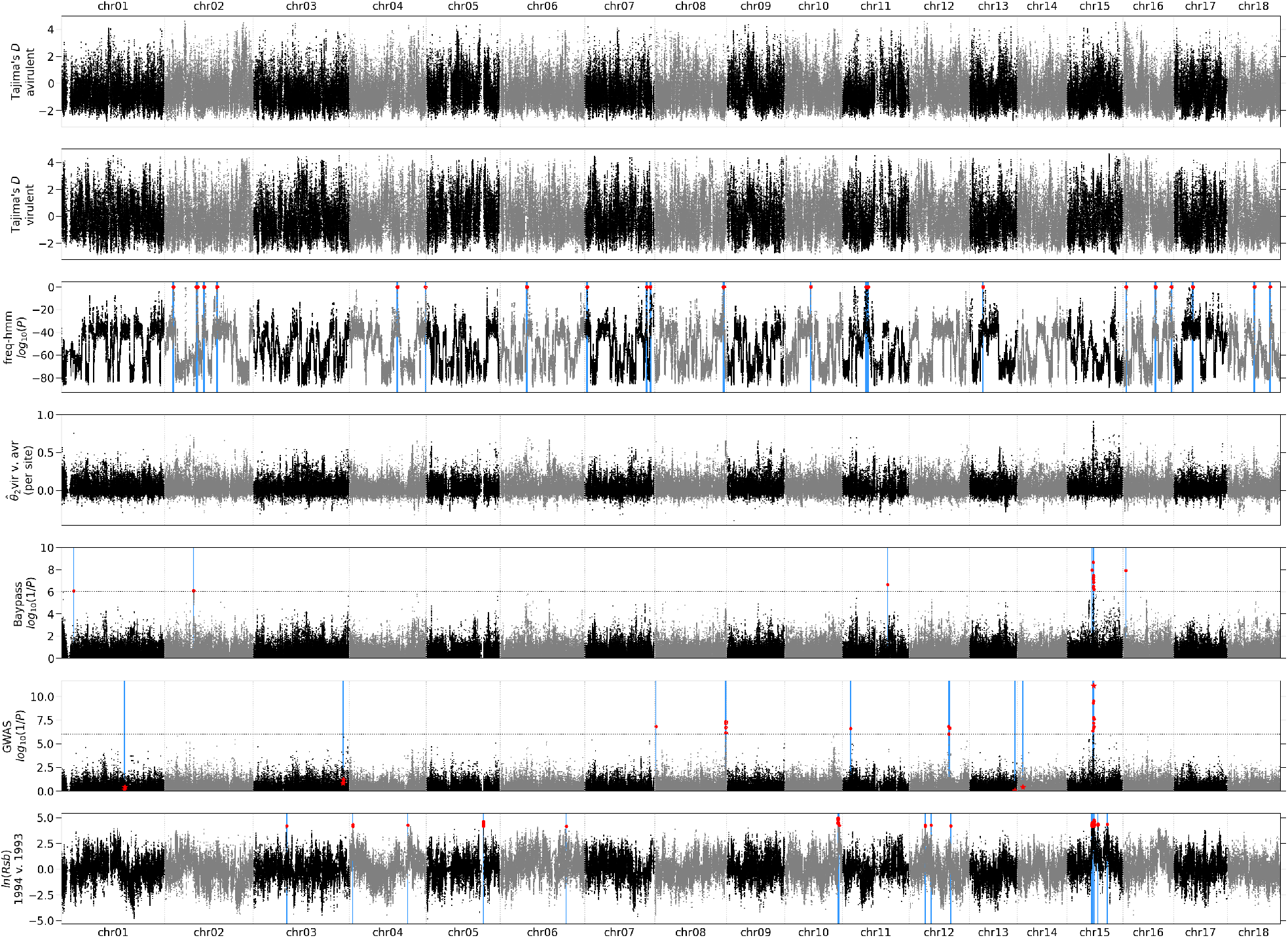
Selective sweep signatures in the *Melampsora larici-populina* genome. Tajima’s *D* and Weir and Cockerham’s 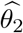 are computed over 1000-bp overlapping windows (step: 250 bp). Each value is placed at the window midpoint. For selective sweep detection methods, the test statistic is given for all SNPs. Significant or outlier values are indicated by larger red disks and their position is outlined by a blue vertical line. For the GWAS results, the SNPs which are significant as cofactors are indicated by a star and the SNPs only passing the threshold in the model without cofactors by red disks.

### Differentiation

For addressing signatures of adaptive differentiation between avirulent and virulent isolates, we screened the variation of Weir and Cockerham’s 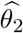 (analogous to *F*_ST_) computed per site. 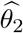 measures the differentiation between the avirulent (1993 and 2008 samples) and virulent (1994 and 1998) clusters. 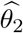 varies widely along the genome with clear peaks at different locations, the highest being in chr15 (position 2,198,745, = 0.92; Fig. 2, fourth panel). A comparison with other differentiation statistics (Additional file 7: Fig. S4) shows that the region with the highest differentiation varies according to the statistic, with for example Jost’s *D* and Hedrick’s 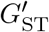 pointing to a region near position 2.61 Mbp of chr09. It should be noted that 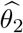 is the only statistic that separates the effect for divergence between virulent and avirulent clusters from the effect of divergence between samples within this clusters.

To screen for potential loci involved in adaptive divergence between virulent and avirulent isolates, we used a Bayesian method aiming to detect adaptive divergence based on allele frequency differentiation between samples (Bay-Pass). The BayPass package [38] provides two models. The core model is based on the sole XtX statistic, which is an analogous to *F*_ST_ [39]. We identified 17 candidate SNPs, of which 12 are located on chr15 (Fig. 2, fifth panel), including the most significant (located at position 2,198,745), and 9 of the 10 most significant, with positions ranging from 2,103,540 to 2,253,676 (Additional file 8: Table S5). The second model of BayPass (covariate model) allows to incorporate phenotypic information. We set the samples 1993 and 2008 as avirulent and the samples 1994 and 1998 as virulent. The results are nearly identical, with 16 candidate SNPs of which 15 (including the 12 on chr15) being also detected by the core model.

### Association genetics

We performed a genome-wide association study (GWAS) to identify SNPs significantly associated with virulence. We used a method that efficiently incorporates loci of large effect as cofactors and increases the power of detection for small effect loci [40]. The best model included seven SNPs as cofactors which altogether explained all the phenotypic variance (Fig. 2, sixth panel, Additional file 9: Fig. S5b). The first cofactor included in the model corresponds to the SNP at position 2,237,229 on chr15 which was the most significant in the initial GWAS scan (Fig. 2, sixth panel). This SNP alone captures 69% of the phenotypic variance (Additional file 9: Fig. S5b), and when treated as a cofactor, all the signal in the region dropped below significance level, suggesting that there is not more than a single potential causal variant in this region. We consider the seven above-mentioned SNPs as candidates, as well as 53 other SNPs that pass the 10^−6^ threshold in the model without cofactors (Additional file 8: Table S5).

### Extended homozygosity

Finally, we investigated extended haplotype homozygosity (EHH), which identifies long-ranging tracts of homozygosity as a potential signature of recent positive selection [41]. Since our data are unphased, we used a variant based on heterozygosity of isolates [42]. We computed iEG (integrated EHH) for all sites of the genome separately for the four samples (Additional file 10: Fig. S6), as well as the ln(*Rsb*) statistic which characterizes the excess of long-range haplotypes of one population with respect to another [42]. We compared sample 1994 to sample 1993, because EHH is meant to detect recent or even on-going selective sweeps, and applied an arbitrary threshold to pick a proportion of 10^−4^ of SNPs. This approach yields 101 candidate SNPs, 61 of them belonging to chr15 and 57 being restricted to the region between 2 and 2.3 Mbp of chr15 (Fig. 2, seventh panel, Additional file 10: Table S6).

### Congruence between genome scan methods

A comparison of the SNPs found using the different methods shows that, of the 15 SNPs which are shared between the two models of BayPass, only three located between 2.2 and 2.4 Mbp on chr15 are shared with GWAS, including the top GWAS candidate, at position 2,237,229 of chr15) and none with the other methods. However, this position is within the region (between 2 and 2.3 Mbp) containing the majority of EHH candidates. Based on BayPass, GWAS and EHH results, we focused on the region of chr15 between positions 2.10 Mbp and 2.30 Mbp (Fig. 3). Taken together, BayPass and GWAS candidates are clustered in two regions: in an intergenic region near position 2,198,500 which is upstream of both flanking genes (`jgi.p|Mellp2_3|84583` and `jgi.p|Mellp2_3|84584`) and in another region spanning three genes (`jgi.p|Mellp2_3|1427900`, `jgi.p|Mellp2_3|71388` and `jgi.p|Mellp2_3|104811`). Of these SNPs, three fall in gene `jgi.p|Mellp2_3|1427900` (positions 2,232,600, 2,233,055 and 2,233,369). The former falls in the 3’ UTR, the second in an intron and the latter causes a non-synonymous change. The top GWAS candidate is located in an intergenic region at position 2,237,229.

**Figure 3:**
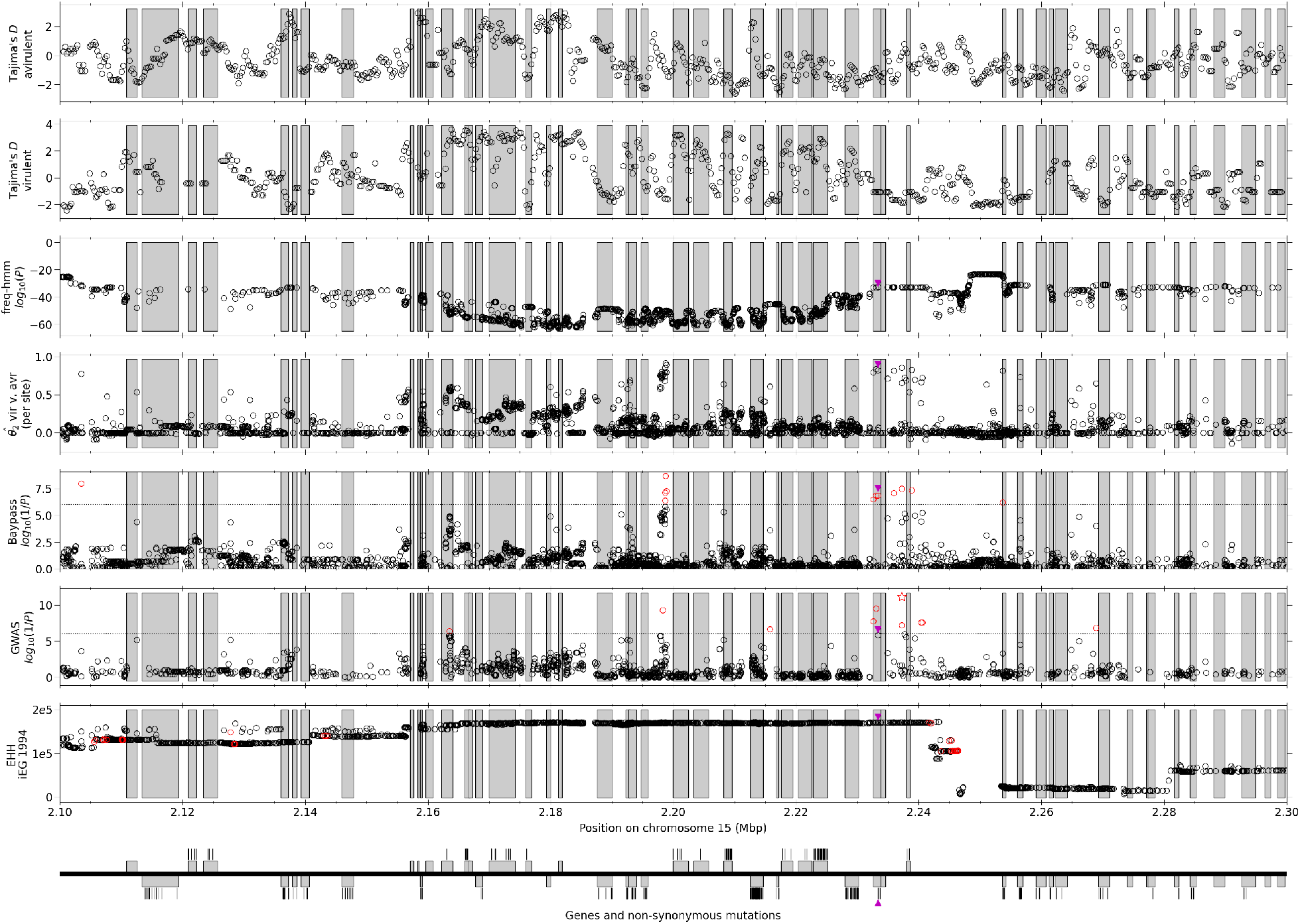
Focus on the candidate region. For all statistics, all genes (full genic region) are indicated by grey frames. The significance threshold is indicated by a dotted line for BayPass and the GWAS. GWAS *P* values from step 1 (without cofactor) are given. Significant SNPs are denoted by red circles (for EHH, significance is assessed by the ln(*Rsb*) ratio of iEG 1994 to iEG 1993) and a red star for the top GWAS SNP. The bottom panel replicates the gene localization (upper frames are forward genes and lower frames are reverse genes). Bars show non-synonymous mutations found in the whole dataset. The one non-synonymous mutation found to be significant in any test is denoted by a purple triangle on all panels.

The EHH analysis shows that the iEG in sample 1994 is consistently high from position 2.16 to 2.24 (including both regions mentioned above) and decreases rather progressively on the left side and more abruptly on the right side near position 2.25 Mbp. The analysis of iEG for the 1998 sample shows a marked decrease of EHH, with a reduced plateau which excludes the `jgi.p|Mellp2_3|84583` gene (Additional file 11: Fig. S7). However, we note that the bottleneck is probably completed in 1998 and that recombination may already have a stronger impact at this point, thereby starting to erode long distance homozygosity. In particular, there is a potential recombination hotspot around position 2.25 Mbp, where iEG values are consistently lower.

### Candidate gene

Interestingly, the three SNPs falling in gene `jgi.p|Mellp2_3|1427900` and the top GWAS candidate are in strong linkage disequilibrium (Additional file 12: Table S6). Except for missing data, all 1994 and 1998 samples have a fixed homozygous genotype for one of the alleles at each SNP except 98GC04 which is heterozygous and 98AB07 which is homozygous for the other allele. 98AB07 is the only avirulent of these two samples, justifying its genotype. Conversely, all 1993 and 2008 samples are either heterozygous or homozygous for the other allele, with the single exception for 08EA95. Therefore, these SNPs, including the one causing a non-synonymous substitution, show a very good correlation with the genotype pattern expected if the virulence was caused by a recessive mutation.

The candidate gene encodes a 219-amino acid protein (ID: 1427900) which is cysteine-rich (seven cysteine residues) and shows no homology to known proteins in *M. larici-populina*, as well as in public databases, including in related Pucciniales species. No N-terminal signal peptide is predicted. Nevertheless the protein is predicted to be non-classically secreted based on SecretomeP 1.0 [43] with a NN score of 0.79 (above the threshold of 0.6). No transmembrane domain was identified with TMHMM v.2.0 [44]. Finally, machine-learning localization prediction tools point to a non-apoplastic protein with signal of chloroplast and nuclear localisations.

## Discussion

Thanks to a sampling design based on collection isolates spanning narrowly the estimated date of the RMlp7 resistance breakdown, we detected consistent molecular signatures pointing to a small region of chr15 and, furthermore, to a specific non-synonymous polymorphism within the gene encoding protein 1427900.

Due to the likely effect of the resistance break-down on the demography, we cannot assume demographic equilibrium for any sample except the 2008 sample and even for this sample the neutrality test statistics suggest that the population is not at equilibrium. In addition, due to the limited availability of collection isolates, the geographical origins of the four samples are heterogeneous, which might cause substructuration of some of the samples. However, substructuration is expected to cause positive values of *F*_IS_ within samples, which does not appear to be the case except for sample 1994. We assume that the strong dispersal abilities and the low inbreeding reproductive system of *M. larici-populina* prevent isolation by distance at a regional scale [45]. Nevertheless, it seemed desirable to search for consistent evidence from complementary methods to conclude about the localization of an avirulence gene candidate.

The molecular signatures of adaptation detected in the current study are of three forms: excess of divergence between virulent and avirulent isolates (tested with BayPass), statistical genotype-phenotype association (tested with GWAS) and presence of long-range homozygosity (tested with EHH). The first two approaches are overlapping since the samples are largely colinear with the virulence status, but use different methodologies. EHH, in contrast, is completely independent and points clearly to the same region. The differentiation and GWAS methods tend to be more precise than EHH and point to one common SNP which is located in an intergenic region. A close survey of the genotypic structure at neighbouring SNPs shows that polymorphism within the gene encoding protein 1427900 (including a non-synonymous mutation) is strongly correlated to the top SNP and that it is, furthermore, consistent with a recessive substitution conferring virulence [46], with all virulent, and no avirulent, isolates being homozygous (with one exception in each case). The non-synonymous substitution can be viewed as the best candidate for determining of the RMlp7 breakdown, although other variants cannot be excluded at this point. Unfortunately, as an obligate biotroph, *M. larici-populina* exhibits features that hinders functional investigations (in particular, no transformation method is available). Further investigations will be needed in order to validate the avirulent character of the candidate gene (e.g. through the use of heterologous systems [47]).

In contrast with the three methods pointing to the candidate region, no selective sweep signatures can be detected in this region through the analysis of the allele frequency spectrum (freqhmm). It seems unlikely that the failure of detecting the region with this kind of method is purely due to a lack a statistical power, because the selective sweep state of the HMM actually decreases in the region. Besides, several other regions in the genome are highlighted at the level of sensitivity we chose and none of them are confirmed by other methods.

Furthermore, we did not find evidence for a strong bottleneck with virulent isolates, especially in the 1994 sample. We expected a strong bottleneck due to the rapidity of the spread of virulence [33, 34]. Bottlenecks cause a transient reduction of diversity followed by a recovery whose rate depends on several factors, including mutation rate and population size [48]. The 1994 sample shows a reduction of diversity compared with 1998, consistent with a bottleneck that would have recovered already. However the substantial level of diversity in 1994 (with values comparable to the other samples), as well as virulence profiles (three different pathotypes in 1994) show that the bottleneck was in fact of weak or moderate intensity. Alternatively, the bottleneck could be actually older than what we envision, which would explain why the bottleneck signatures are relatively weak. However, due to active surveillance of rust disease on cultivated poplars [31] it is rather unlikely that a large population of virulent *M. larici-populina* existed for a significant amount of time without being detected.

The most likely explanation of the low impact of the bottleneck on the allele frequency spectrum and of the low reduction of diversity in the virulent genetic group is a soft selective sweep. Soft selective sweeps occur when adaptation is mediated by a mutation which was actually segregating in the original population and are thought to be more likely to be involved in fast evolution [49]. The extent to which soft selective sweeps contribute to adaptive evolution is debated [50, 51, 52]. In our study, a soft selective sweep appears as the most likely hypothesis, consistently with the fact that adaptation to human-introduced host resistance was prone to trigger fast adaptation by the pathogen. Consistently with this hypothesis, the putatively causal allele can be found in a heterozygous state in the 1993 population which predates the putative breakdown event. In addition to a soft selective sweep, the life history of *M. larici-populina* reduces inbreeding due to mating types, the need of two hosts to complete the biological cycle and high dispersal abilities [53]. These features likely reduced further the extent of hitch-hiking around the putative avirulence locus. In particular, this can explain why the signatures of the selective sweep do not expand further than 0.2 Mbp from the putative selected locus (Fig. 3) and the lack of strong signatures on pairwise linkage disequilibrium on chr15 (Additional file 6: Fig. S3).

Plant pathogen effectors are classically searched on the basis of common features such as secretion signals, small size (< 300 amino acids), high cysteine content and lack of homology in other species [54]. The candidate gene found in this study shares some but not all these features. In particular, the lack of a classical secretion signal would have excluded this gene from screening analysis based on common features such as those already performed on *M. larici-populina* [53, 55].

We cannot exclude that the true causal mutation is elsewhere in the same genomic region. The non-synonymous substitution is the best candidate *a priori*, but silent mutations can have an effect as well, such as mutations in regulatory elements [56]. Furthermore, our analysis is based on the current annotation of the genome of the reference isolate of *M. larici-populina* (98AG31) which is actually a virulent individual. It is possible that the annotation of the avirulence gene is incorrect in the virulent reference isolate because of a frame-shifting mutation or that a large deletion occurred. *De novo* genome assembly of an avirulent isolate using a long read sequencing approach would be required to exclude these possibilities.

## Conclusion

Our study identified the best candidate avirulence gene in *Melampsora larici-populina* so far, based on the survey of genomic diversity. By considering samples framing narrowly the supposed date of the resistance breakdown, we maximized power to identify the causal locus. Our study illustrates the benefit of monitoring the diversity of pathogens through collections of living samples, in order to trace back any significant evolutionary transition that may occur. In complement, we combined a set of complementary genome scan methods to increase accuracy of the screen for adaptation. Our study suggests that features of the pathogen life-cycle such as sexual reproduction, high dispersal, and high levels of diversity foster fast adaptation while resulting in a genetically diverse virulent population which has lost virtually none of its potential for further adaptation.

## Methods

### Fungal material

Four samples were designed based on the population structure previously described in [34] (Additional file 1: Table S1). We extracted isolates from a collection available in the laboratory. To ensure their purity, genotyping was performed using 25 microsatellite markers with the method used in [34]. Virulence profiles were characterized on a differential set of nine poplar genotypes each carrying a single qualitative resistance to *Melampsora larici-populina* (RMlp1: *Populus* × *euramericana* ‘Ogy’; RMlp2: *P*. × *jackii* ‘Aurora’; RMlp3: *P*. × *euramericana* ‘Brabantica’; RMlp4: *P*. × *interamericana* ‘Unal’; RMlp5: *P*. × *interamericana* ‘Rap’; RMlp6: *P*. × *interamericana* ‘84B09’; RMlp7: *P*. × *interamericana* ‘Beaupré’; RMlp8: *P*. × *interamericana* ‘Hoogvorst’; RMlp9: *P. deltoides* ‘L270-3’) and on the universal susceptible cultivar *P*. × *euramericana* ‘Robusta’ as positive control. Poplar plants were grown during 2-3 months in a glasshouse at 20-24°C, with a 16 hours photoperiod, as previously described in [57]. We excised 12-mm discs on leaves (index 7 to 14) and placed them in flotation on deionized water in 24-well polystyrene cell culture plates, abaxial surface up. A suspension (approximately 3.2×10^6^ spores/mL) of each strain was deposited as 1-µl droplets on each disc. Culture plates were incubated for 13 days at 19 ± 1°C under continuous illumination before scoring. Isolates resulting in at least ten pathogenic lesions per leaf disk were scored as virulent with respect to the corresponding resistance. Three replications were performed.

### DNA isolation

The DNA isolation method was performed on 50 mg of urediniospores as previously decribed in [36, 57]. Quality and quantity of recovered high molecular weight DNA was assessed by electrophoresis on agarose gel, by spectrophotometry (Nanodrop, Saint-Remy-lés-Chevreuse, France) and with the QuBit fluorometric quantitation system (Life Technologies, Villebon-sur-Yvette, France).

### Genome re-sequencing, filtering and mapping

DNA was used for sequencing by Beckman Coulter Genomics (Grenoble, France) for 22 isolates and at the Joint Genome Institute for the remaining 54 (Additional file 1: Table S1). Each library was quantified by qPCR and sequenced on the Illumina HiSeq2000 platform as paired-end 150 bp reads.

The reads were mapped on the *M. larici-populina* reference genome v2.0, available via the JGI fungal genome portal MycoCosm [58]. The assembled genome contains 18 chromosomes (101.4 Mbp) and 493 unmapped scaffolds (8.4 Mbp). The unmapped scaffolds, representing 8% of the total genome length (average length: 17 Kbp, N50: 36 Kbp), were not considered. All reads were aligned on the reference genome using the bwa software version 0.7.13 [59]. We fixed the maximum number of differences for each read against the reference to 2. All others options were used with default values. We used SAMtools version 1.3 [60] with default parameters to compute the number of mapped reads, perform genotype calling and export variant call (VCF) format files.

SNPs were filtered using EggLib version 3 [61]. Genotypes were assigned based on the difference of Phred-scaled likelihoods (PL values in VCF files) between the best genotypes and all others as implemented in EggLib (`threshold_PL` = 30). Genotypes with a depth below 20 or above 200 reads were called as missing to exclude sites falling potentially in repetitive DNA. After genotype calling for all samples, we excluded sites with more than two alleles overall and less than 10 non-missing isolates in any of the four samples.

### Phylogenetic tree and population structure

To build a phylogenetic tree of all isolates, we computed the pairwise distance based on all SNPs. Pairwise distances were computed as the rate of pairwise genotypic differences for a random subset of sites without missing data for the two considered isolates, without correction. One percent of sites was drawn independently for all pairs of individuals. The tree was reconstructed using Phylip v 3.696 [62] using the neighbour-joining method. For representation we used the Interactive Tree of Life online editor [63]. The population structure of the isolates was assessed with DAPC implemented in the Adegenet package in R [37]. A random subset of 10% of sites was used. The number of genetic groups was assessed based on decreasing Bayesian information criterion (BIC) values.

### Analysis of polymorphism

Linkage disequilibrium (LD) was computed as *r*^2^ using EggLib. Pairwise genotypic LD was computed separately for all four samples and for all pairs of sites of the same chromosome with a distance at most 1 Mbp, excluding sites with a genotype at a frequency above 75% or an overall frequency of missing data above 10%. To smooth LD decay curves in the graphical representation, we computed quantiles (including the median) of *r*^2^ values grouped based on windows of 1000 bp of pairwise distance. We also monitored the window where the average *r*^2^ passed below different thresholds and the average *r*^2^ at given distance points. We also computed pairwise genotypic LD between all pairs of SNPs of each chromosome (also separately for the four samples), considering at most one SNP per Kbp of the reference genome.

We computed summary statistics using EggLib either for individual SNPs, in sliding windows along the chromosomes or for the whole genome (considering the whole set of sites). We defined window boundaries on the reference genome and processed complete windows only. For the sake of symmetry, we shifted window starting points in order to leave equal stretches of unanalysed sequence at both ends of each chromosome. All per-site statistics were expressed relatively to the number of considered sites (variable or not) within the considered window (or overall), ignoring sites excluded due to an excess of alleles or missing data. Within-population statistics were computed within the four samples and for the whole dataset. Differentiation statistics were computed for the whole datasets (four groups) and for pairwise comparisons. We measured the divergence between virulent and avirulent isolates by assessing the between-clusters differentiation using Weir and Cockerham’s method where clusters were defined as the 1993 and 2008 samples (avirulent) on one hand and the 1994 and 1998 samples (virulent) on the other hand. Alternatively, we computed differentiation statistics considering two populations with respectively virulent and avirulent isolates. In the latter case, based on phenotyping results, the 98AB07 isolate was treated as avirulent (Additional file 1: Table S1).

### Genome scan methods

BayPass [38] version 2.1 was used with default parameters. For the covariate model, virulence status was encoded as an environmental value with a constant value −1 for avirulent populations and +1 for virulent populations. All polymorphic sites were used. The significant threshold was set to 10^−6^.

EHH was estimated for all polymorphic sites with EggLib, separately for all four samples (note that not all sites were polymorphic in all samples). The main statistic is iEG which integrates the site-wise EHHS statistic [42]. Each site was treated in turn as the core site and iEG was computed by integrating in both directions, until EHHS reached a threshold of 0.2 (or the chromosome limit was reached). To identify candidates, we computed ln(*Rsb*) as described in [42] and selected all loci with a ln(*Rsb*) in the right tail of the distribution (probability density 10^−4^).

GWAS was performed using a mixed-model approach including multiple loci [40], considering the virulence as phenotype and the genotypes as alleles. Since missing data are not supported, they were imputed by assigning the majority genotype to isolates with missing data. Only sites with a minor allele frequency of at least 0.05 were considered. To incorporate the information of genetic structure (including the differentiation between the four samples), we computed a kinship matrix [64] which was included in the model as the covariance matrix for a random polygenic effect. We used the mBonf and EBIC criteria to select the best model from the forward-backward search [40]. We report *P* values from the initial GWAS scan (no cofactor). Candidate loci are determined as those identified as cofactors in the best multi-locus model and/or which are significant at a 10^−6^ threshold in the initial scan.

The freq-hmm method [65] was applied to all polymorphic sites on allele frequencies using the folded spectrum mode, independently for all four samples. The starting value for *θ* was set to 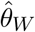 as computed for the whole dataset for each sample. The analysis was performed separately for all chromosomes and the *k* parameter (which controls the detection rate) was set to 10^−20^.

### Candidate gene analysis

The encoded protein was evaluated by comparison to proteins of known function in the nonredundant GenBank database. Signal peptide prediction in amino acid sequences was performed using SignalP 5.0 [66]. Non-classical secretion was predicted using SecretomeP 1.0 [43] with the normal threshold of neural network output score of 0.6 as recommended by the authors. Transmembrane domain detection was performed with TMHMM v.2.0 [44] and PHOBIUS [67]. Potential subcellular localization was predicted in silico using online tools APOPLASTP [68] and LOCALIZER [69].

## Acknowledgements

We are grateful to Jérémy Pétrowski for the production of biological material for our experiments. We thank Pierre Gladieux, Martin Lascoux, Christophe Lemaire and Mathieu Siol for comments on earlier versions of the manuscript.

## Availability of data and materials

Sequencing reads for isolates sequenced by Beckman Coulter Genomics have been deposited in Genbank’s Sequence Read Archive (https://www.ncbi.nlm.nih.gov/sra). Sequencing reads for isolates sequenced by the JGI are available on the JGI Genome Portal under Proposal ID 1416 (doi: 10.25585/1488093). Accession numbers to the SRA and JGI Project IDs, respectively, are given in Additional file 1: Table S1.

All scripts used to perform the analysis of polymorphism and processing of genome scan results, including scripts used to generate figures, are available in a dedicated git repository (https://gitlab.com/demita/mlp-genomicsvir7).

## Authors’ contributions

SD, PF, FH and SDM designed the study. AP, AM and AA performed experiments and generated biological material. AL, KWB, HN, CA and IVG sequenced and analysed part of the isolates. AP and SDM analysed data with contributions of AM, CL and VS. AP, AM, CL, VS, SD, PF, FH and SDM discussed the results. AP and SDM wrote the manuscript with contributions from AM, CL, PF and FH.

## Additional Files

### Additional file 1

**Table S1**. Characteristics of sampled isolates and sequencing results. The pathotype column gives the index of all tested resistances that each sample was able to overcome (tested resistances: RMlp1 to RMlp9). If none, a value of 0 is given. For each isolate, the accession number for sequencing reads is given, either to Genbank’s Sequence Read Archive (SRA) or as a Project ID within the JGI Genome Portal.

### Additional file 2

**Table S2**. Detection of polymorphic sites. The table shows, for all 18 chromosomes, the reference chromosome length, the number of un-called positions, the number of called positions, the number of fixed positions (where only one allele is present, whether or not it is the reference allele) and the number of variable positions. The proportions of uncalled, called, fixed and variable positions are given with respect to the chromosome length and the proportions of fixed and variable positions are given with respect to the number of called positions.

### Additional file 3

**Figure S1**. Result of the DAPC analysis. **a** Selection of the number of groups (‘clusters’ in the figure) using the Bayesian information criterion (BIC). Representation of the number of isolates of each samples assigned to each group in the analysis with **b** two groups, **c** three groups and **d** four groups. Membership coefficients of each isolate in the analysis with **e** two groups, **f** three groups and **g** four groups. In panels **e**-**g**, the four samples are separated by thick lines and the colours identifying groups are arbitrary.

### Additional file 4

**Table S3**. List of statistics computed in this study. **Table S4**. Genome-wide statistics. More details are given in the file.

### Additional file 5

**Figure S2**. Pairwise values of *r*^2^ between all pairs of site. **Table S5**. Linkage disequilibrium decay statistics.

### Additional file 6

**Figure S3**. Pairwise linkage disequilibrium for all chromosomes. Each page of this document contains a half-matrix representing a chromosome. For a given chromosome, one SNP per Kbp has been selected and the genotypic *r*^2^ has been computed for all pairs and is represented on a grayscale (a legend is included in each page). The physical position of SNPs is expressed in Mbp at the bottom of each half-matrix. The analysis has been performed separately for the four samples. If a Kbp window didn’t have any suitable SNP, the row is white.

### Additional file 7

**Figure S4**. Diversity statistics over a sliding window. This figure completes Fig. 2 with additional statistics: 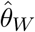 *D* per sample and additional population differentiation statistics, including values computed per site. The panel for 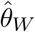 is cropped to a maximal value of 0.01, discarding larger values.

### Additional file 8

**Table S5**. Results of selective sweep detection methods. Spreadsheet providing the list of SNPs passing thresholds for the five selective sweep detection methods. The first sheet gives the list of SNPs significant with at least two methods (discarding SNPs common to the variants of Bay-Pass only). For BayPass core and covariate models the list of SNPs is provided with their individual *P* value. For GWAS, the list of candidate SNPs comprised 54 SNPs passing the threshold in the model without cofactors, and the 7 SNPs declared as cofactors in the best models (their rank is indicated). Only one SNP is in common. For EHH, the value of iEG in the 1994 test and the ln(*Rsb*) for the 1994 to 1993 comparison are given. For freq-hmm, candidate selection sweep regions are specified by their coordinates along with the maximum posterior probability within the region.

### Additional file 9

**Figure S5**. Variance partitioning along the GWAS forward-backward procedure. Each step corresponds to the forward inclusion (until the criterion is reached) and then to the backward elimination of loci as cofactors.

### Additional file 10

**Figure S6**. Results of the EHH analysis. The six first panels represent iEG (integrated haplotypic EHH) along the genome for the four samples and the virulent and avirulent clusters. The last four panels represent the ln(*Rsb*) ratio statistic for pairwise comparisons.

### Additional file 11

**Figure S7**. Details of EHH statistics in the focus region. The figure template corresponds to Fig. 3. The iEG is given for all four samples along with ln(*Rsb*) for three comparisons: 1994 to 1993, 1994 to 2008, and 1998 to 1993. Significant values for 1994 and 1998 iEG are denoted by red circles and are based to the ln(*Rsb*) comparison to 1993.

### Additional file 12

**Table S6**. Genotypes of candidate SNPs on chr15. The genotypes are given in a plain text table (space separated). The top GWAS candidate as well as three nearby SNPs highlighted by BayPass falling within gene `jgi.p|Mellp2_3|1427900` are presented.

