## Supplementary figures and images for "Genomic signatures of a major adaptive event in the pathogenic fungus *Melampsora larici-populina*"

### Additional file 03

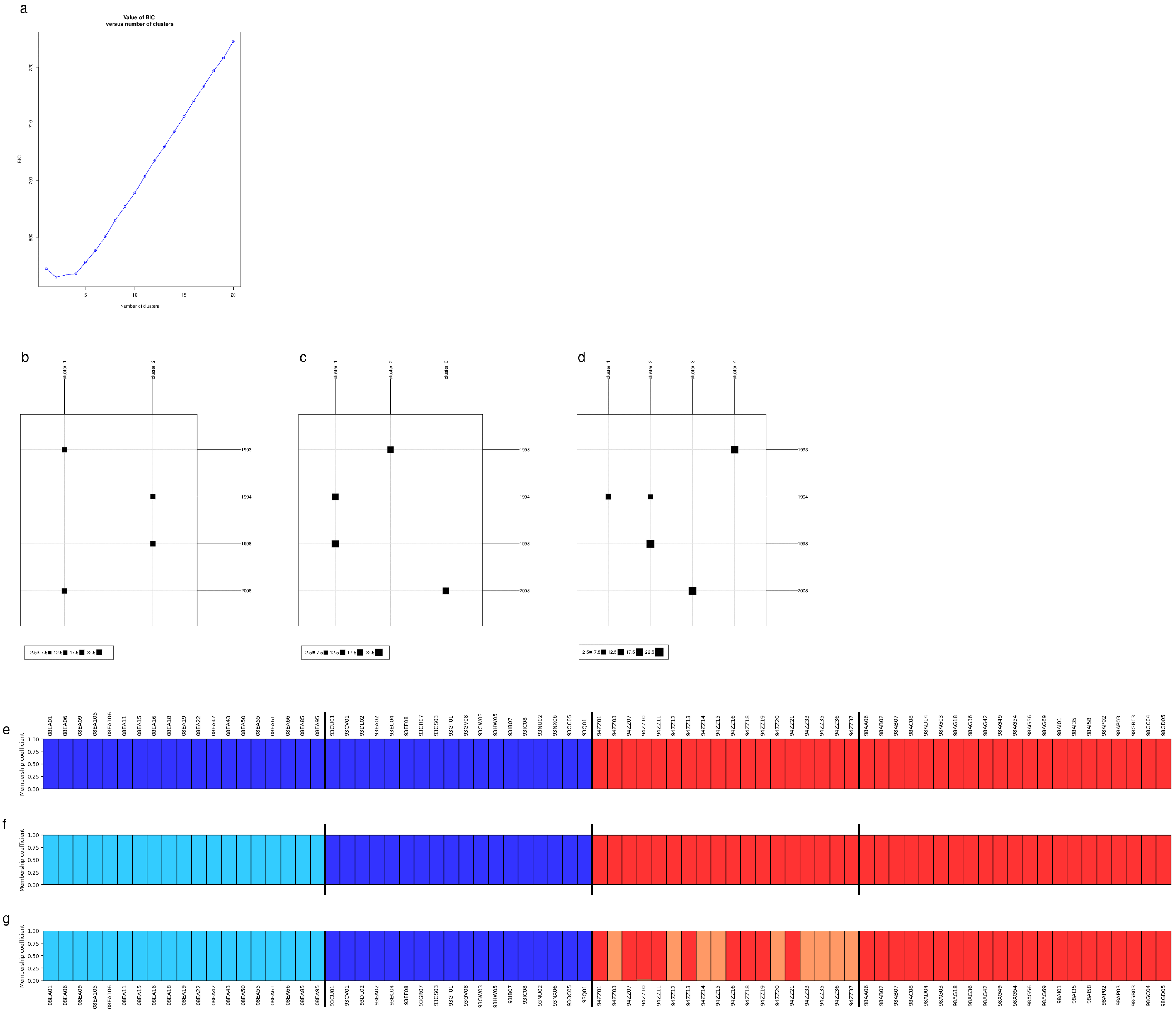

### Additional file 06

#### chr01

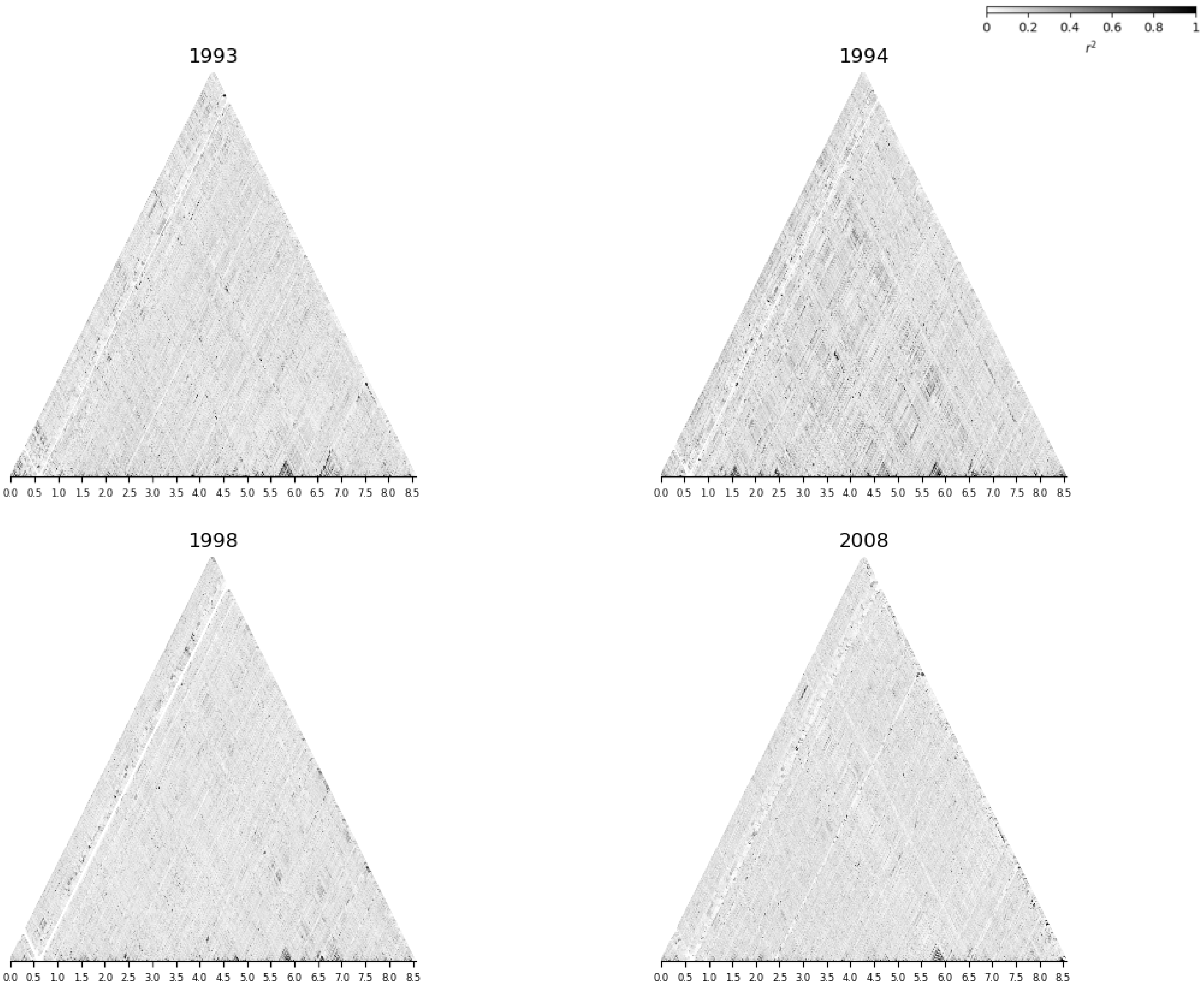
0 0.2 0.4 0.6 0.8 1
r²
1993
1994
1998
2008
0.0 0.5 1.0 1.5 2.0 2.5 3.0 3.5 4.0 4.5 5.0 5.5 6.0 6.5 7.0 7.5 8.0 8.5

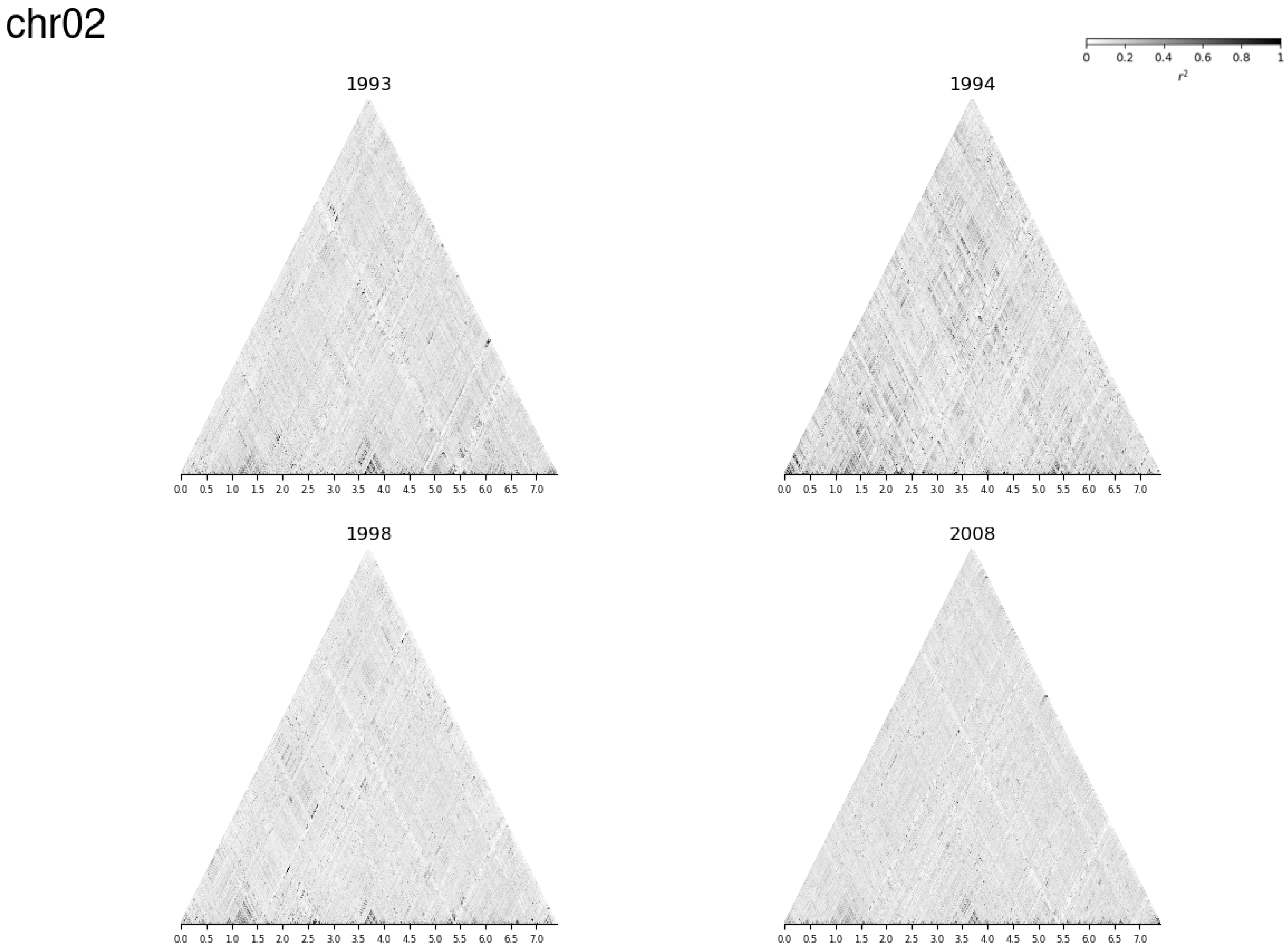

chr02
0
0.2
0.4
0.6
0.8
1
$r^2$
1993
1994
1998
2008
0.0 0.5 1.0 1.5 2.0 2.5 3.0 3.5 4.0 4.5 5.0 5.5 6.0 6.5 7.0

#### chr03

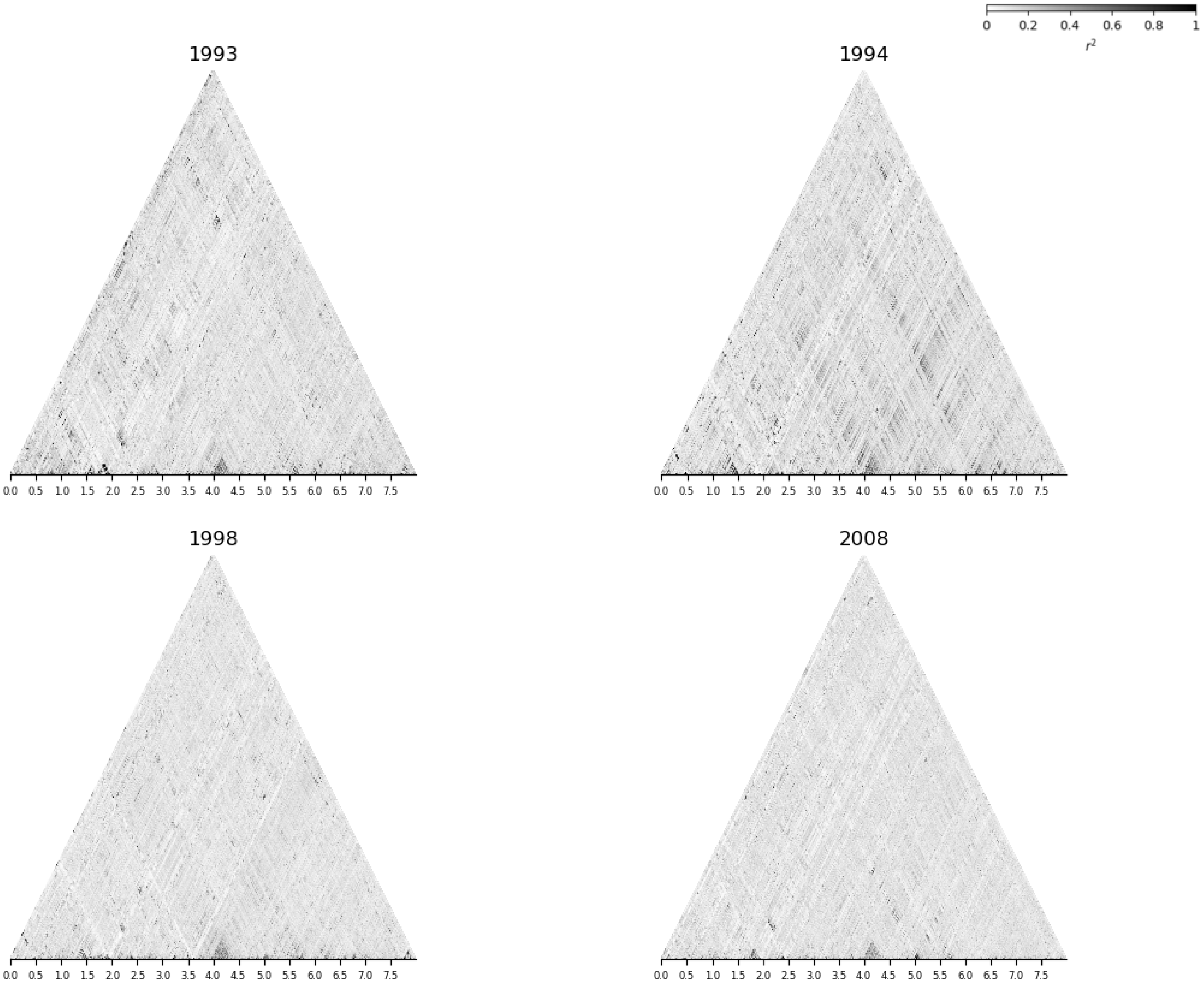

0
0.2
0.4
0.6
0.8
1
r²
1993
1994
1998
2008
0.0 0.5 1.0 1.5 2.0 2.5 3.0 3.5 4.0 4.5 5.0 5.5 6.0 6.5 7.0 7.5

#### chr04

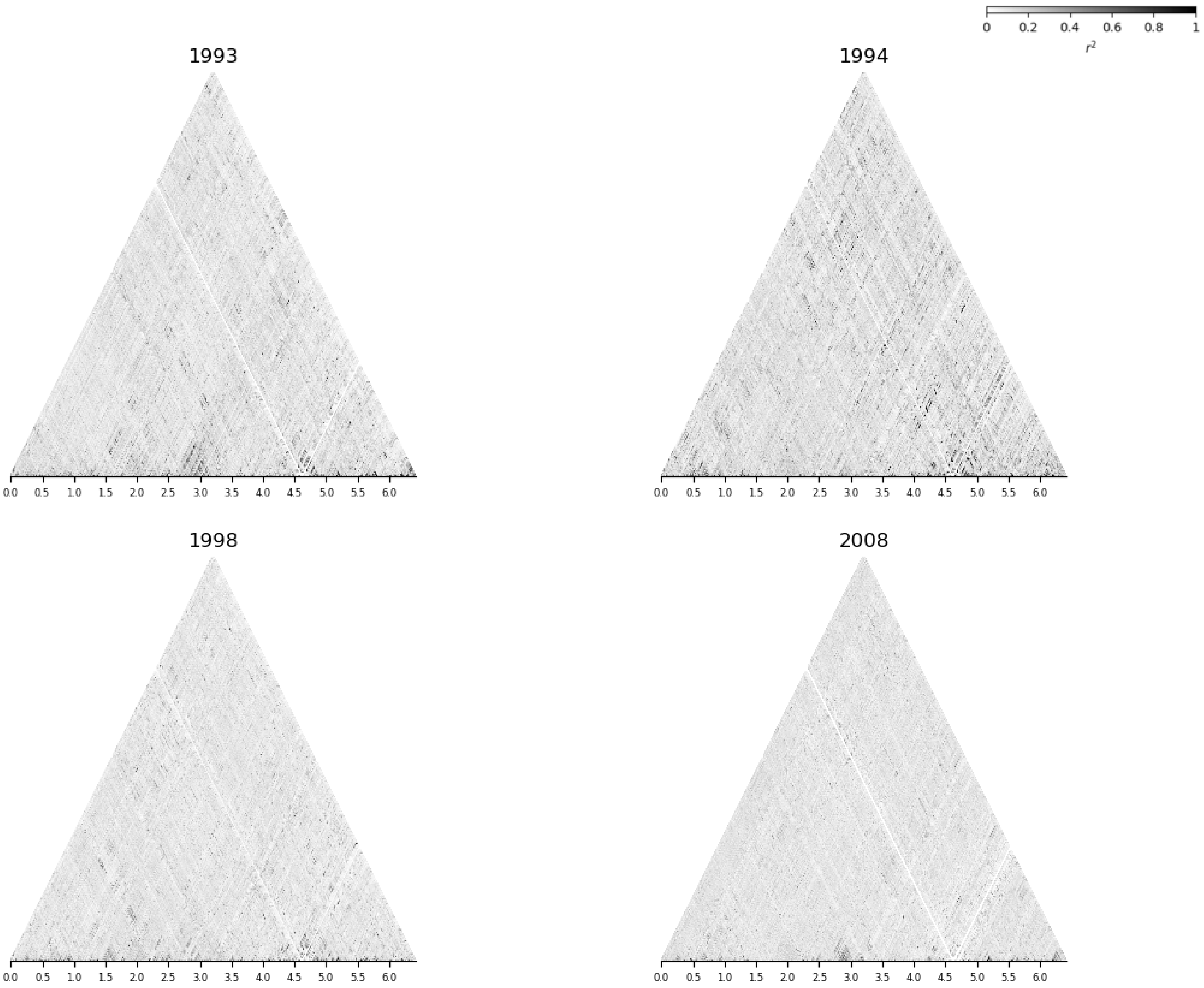

0 0.2 0.4 0.6 0.8 1
$r^2$
1993
1994
1998
2008
0.0 0.5 1.0 1.5 2.0 2.5 3.0 3.5 4.0 4.5 5.0 5.5 6.0

chr05

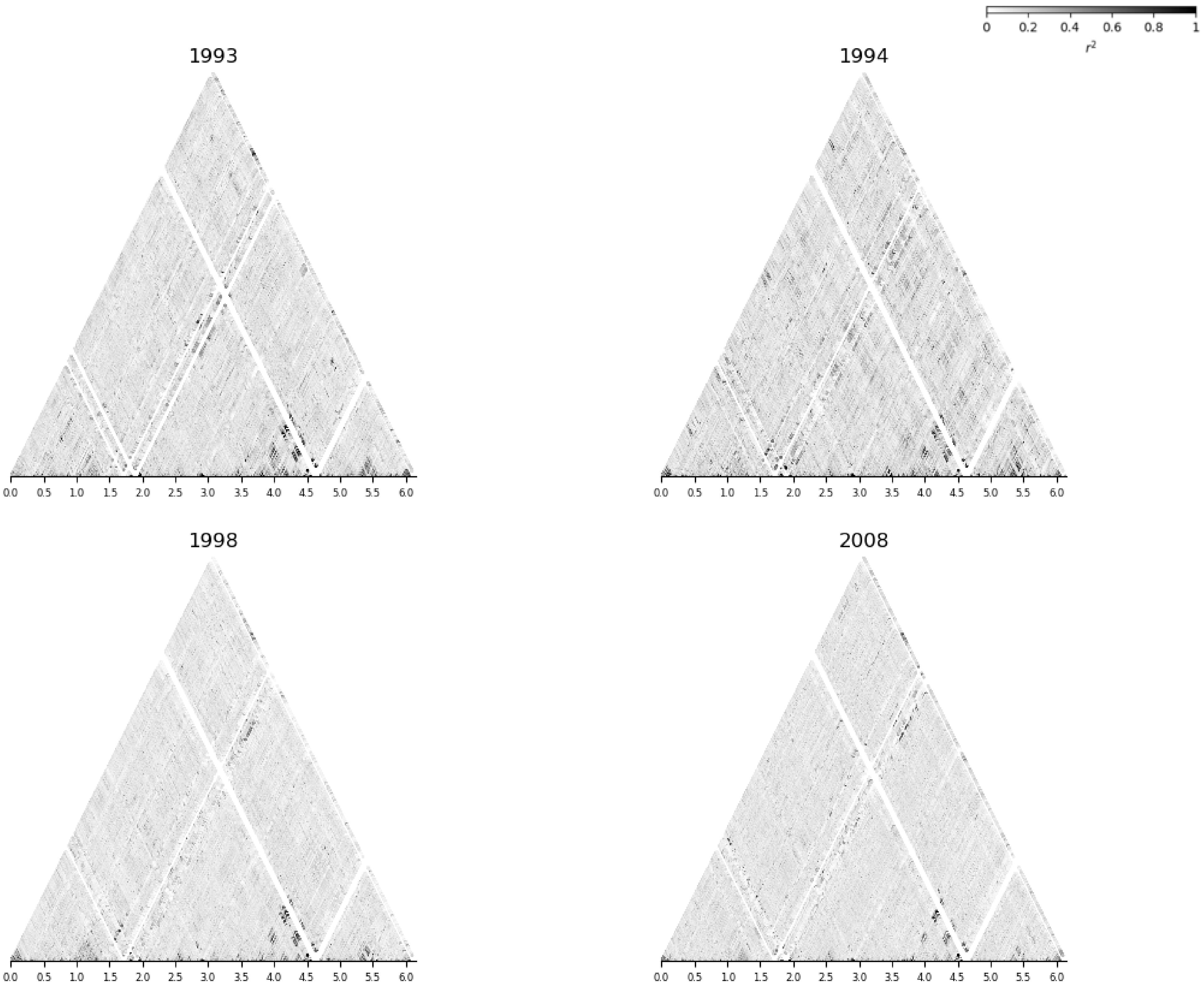

0
0.2
0.4
0.6
0.8
1
$r^2$
1993
1994
1998
2008
0.0
0.5
1.0
1.5
2.0
2.5
3.0
3.5
4.0
4.5
5.0
5.5
6.0

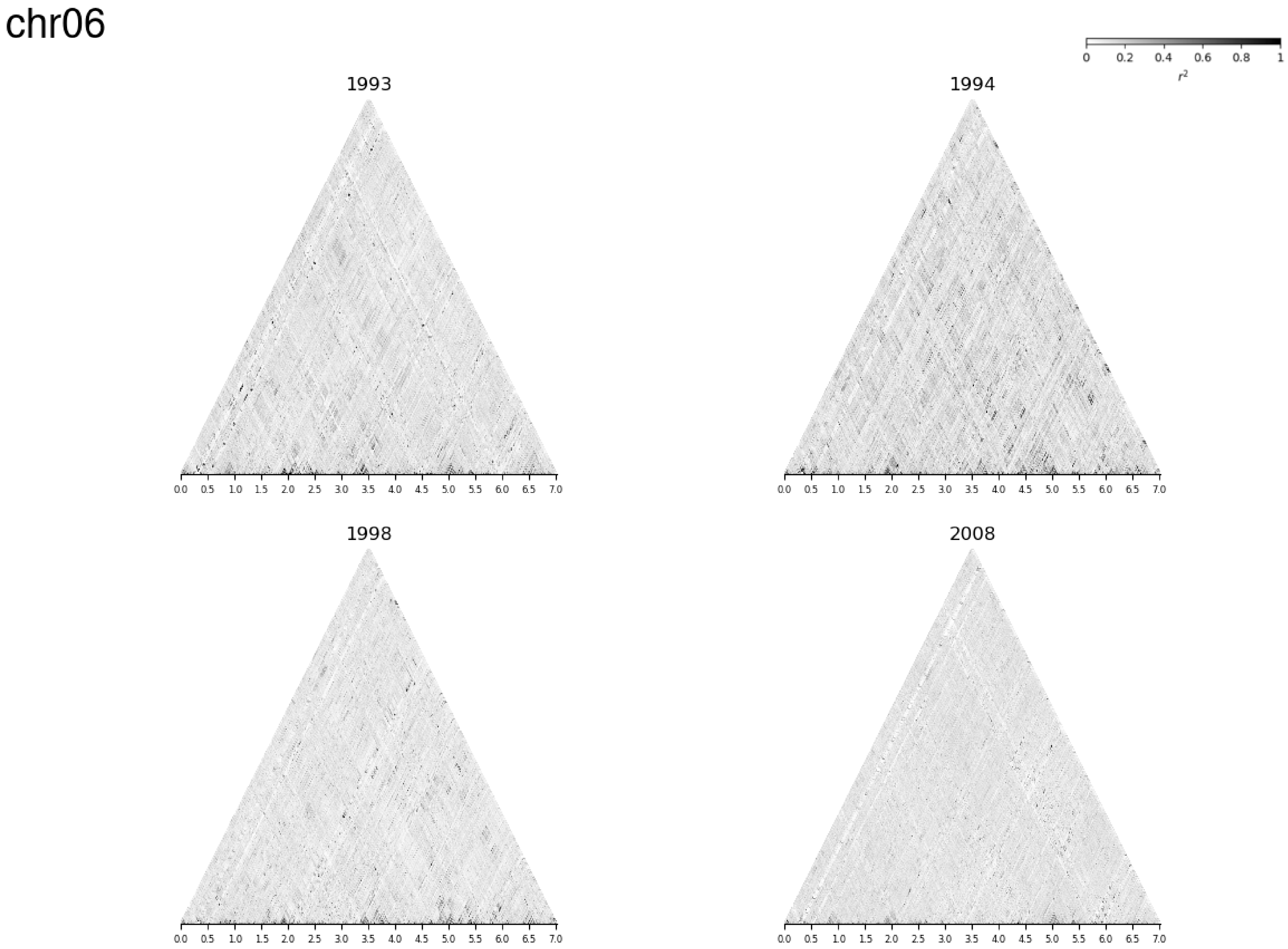
chr06
0
0.2
0.4
0.6
0.8
1
$r^2$
1993
1994
1998
2008
0.0 0.5 1.0 1.5 2.0 2.5 3.0 3.5 4.0 4.5 5.0 5.5 6.0 6.5 7.0

#### chr07

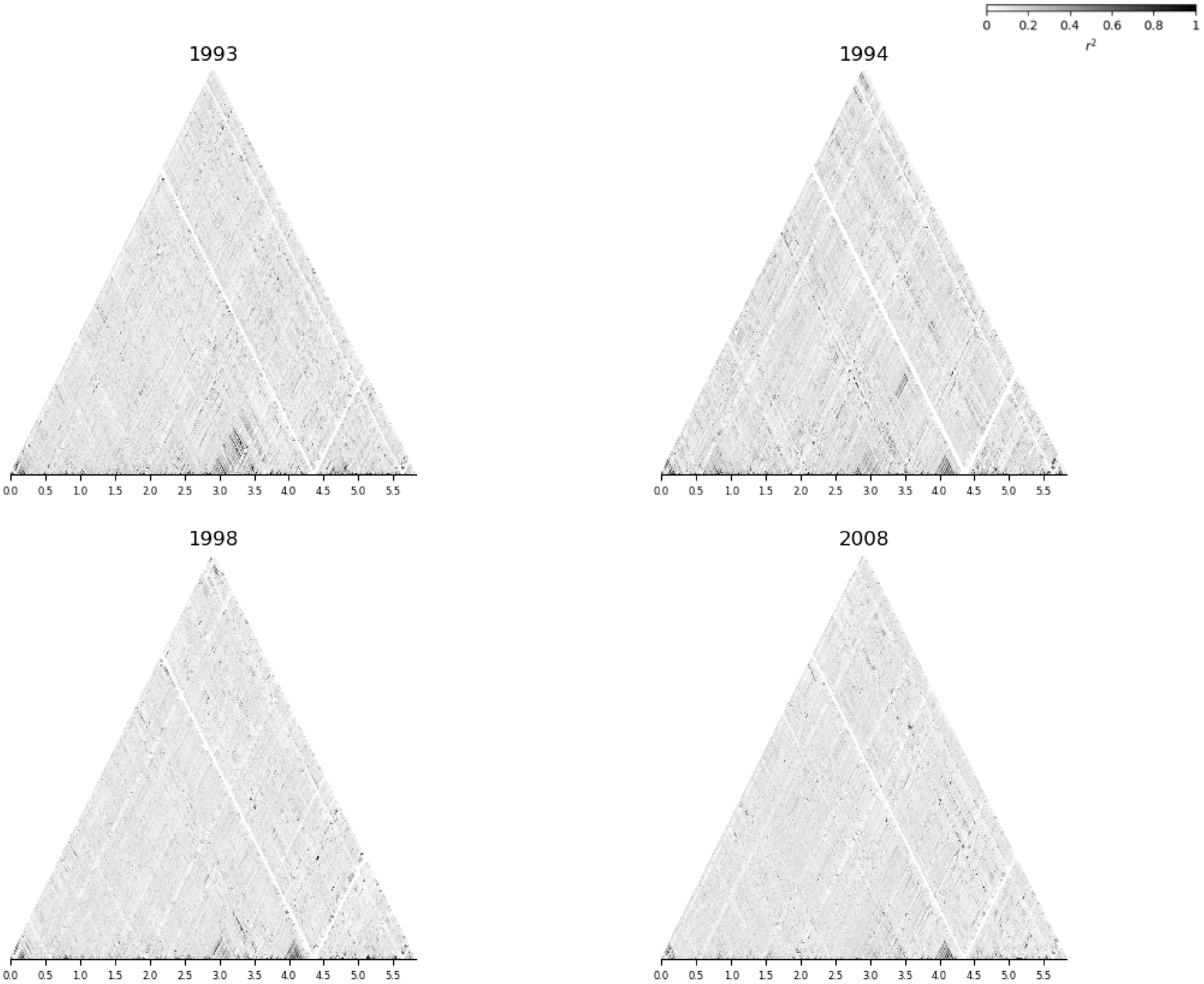
0
0.2
0.4
0.6
0.8
1
$r^2$
1993
1994
1998
2008
0.0
0.5
1.0
1.5
2.0
2.5
3.0
3.5
4.0
4.5
5.0
5.5

#### chr08

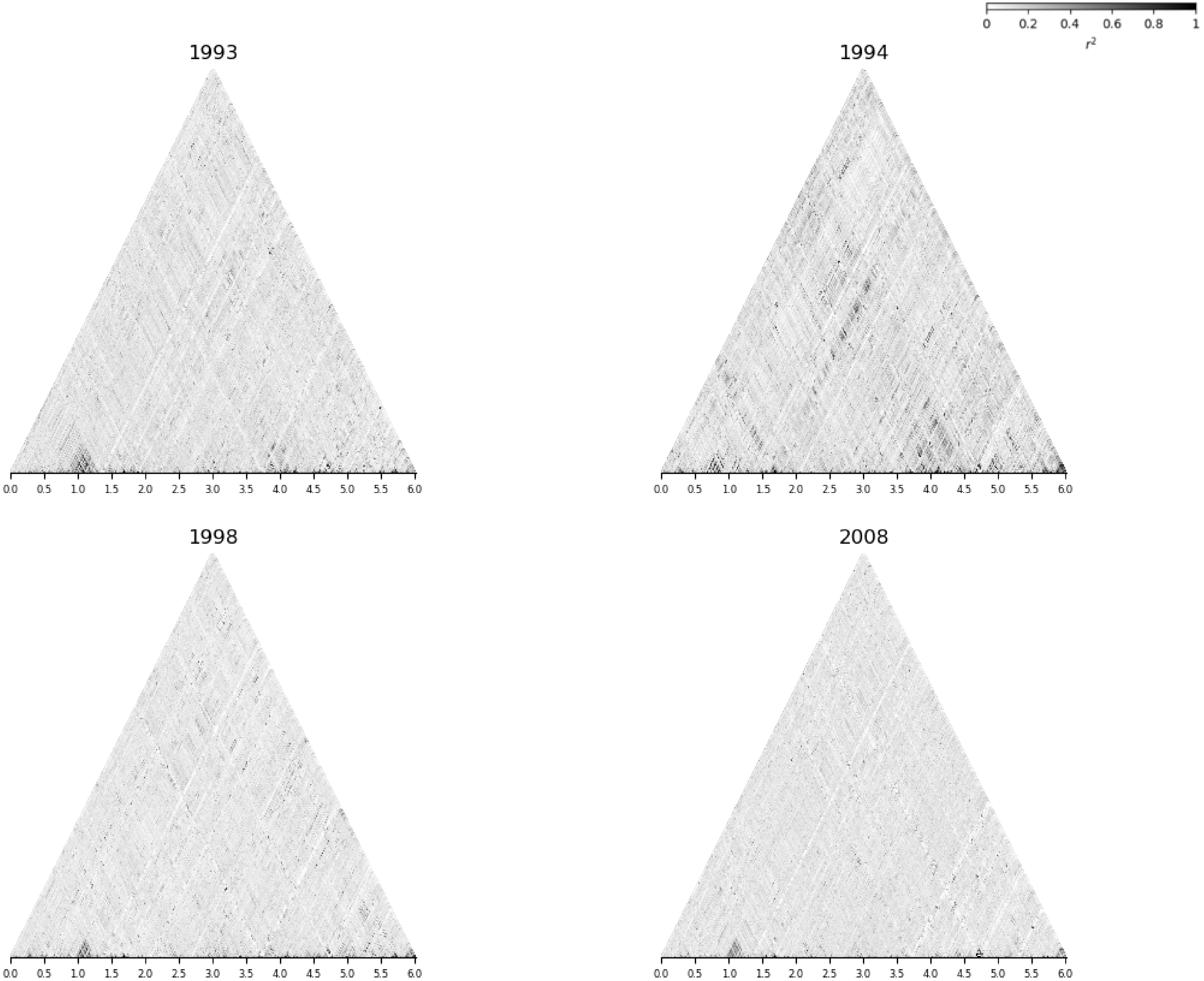

0
0.2
0.4
0.6
0.8
1
$r^2$
1993
0.0 0.5 1.0 1.5 2.0 2.5 3.0 3.5 4.0 4.5 5.0 5.5 6.0
1994
0.0 0.5 1.0 1.5 2.0 2.5 3.0 3.5 4.0 4.5 5.0 5.5 6.0
1998
0.0 0.5 1.0 1.5 2.0 2.5 3.0 3.5 4.0 4.5 5.0 5.5 6.0
2008
0.0 0.5 1.0 1.5 2.0 2.5 3.0 3.5 4.0 4.5 5.0 5.5 6.0

#### chr09

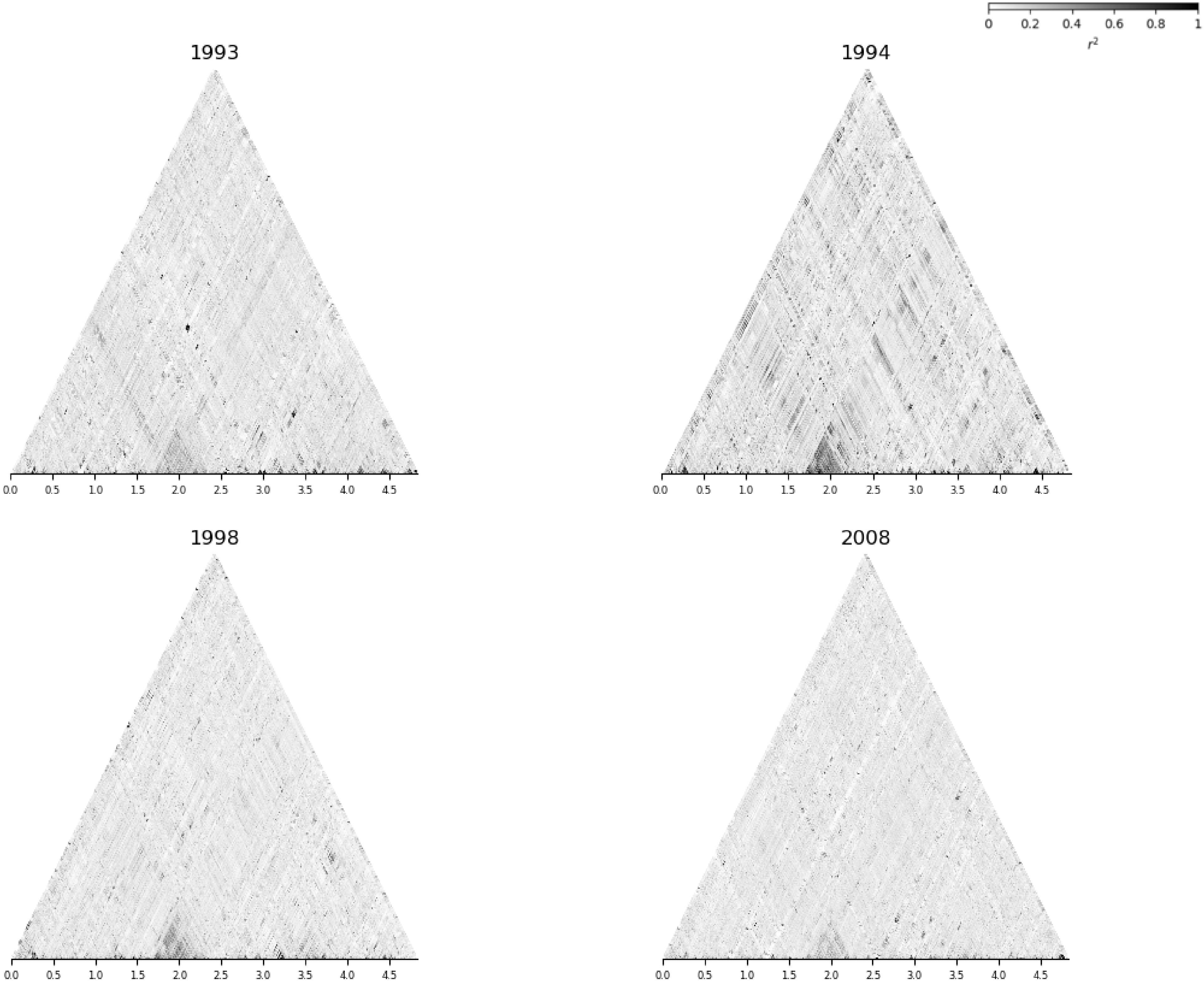

chr10

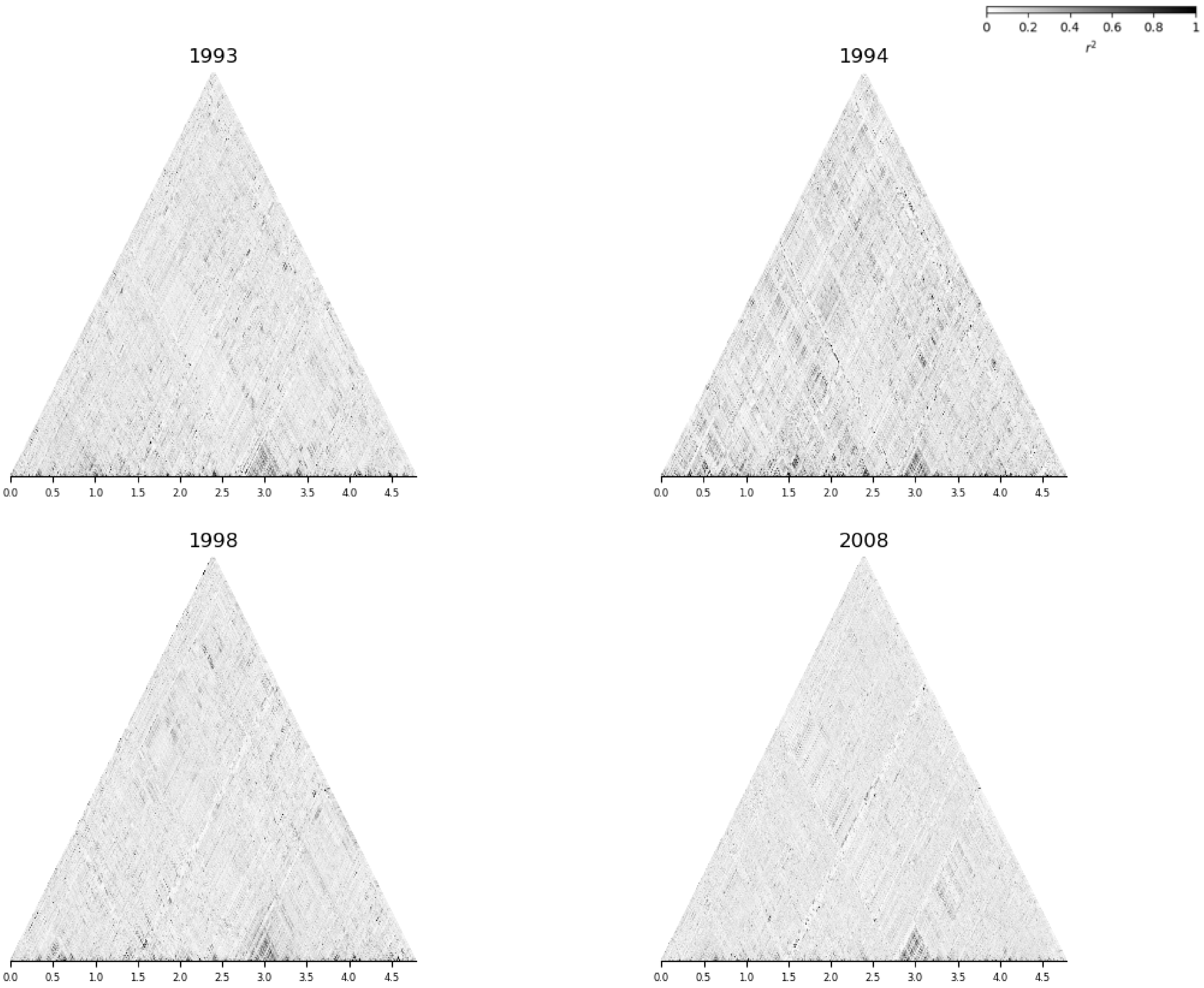

chr11

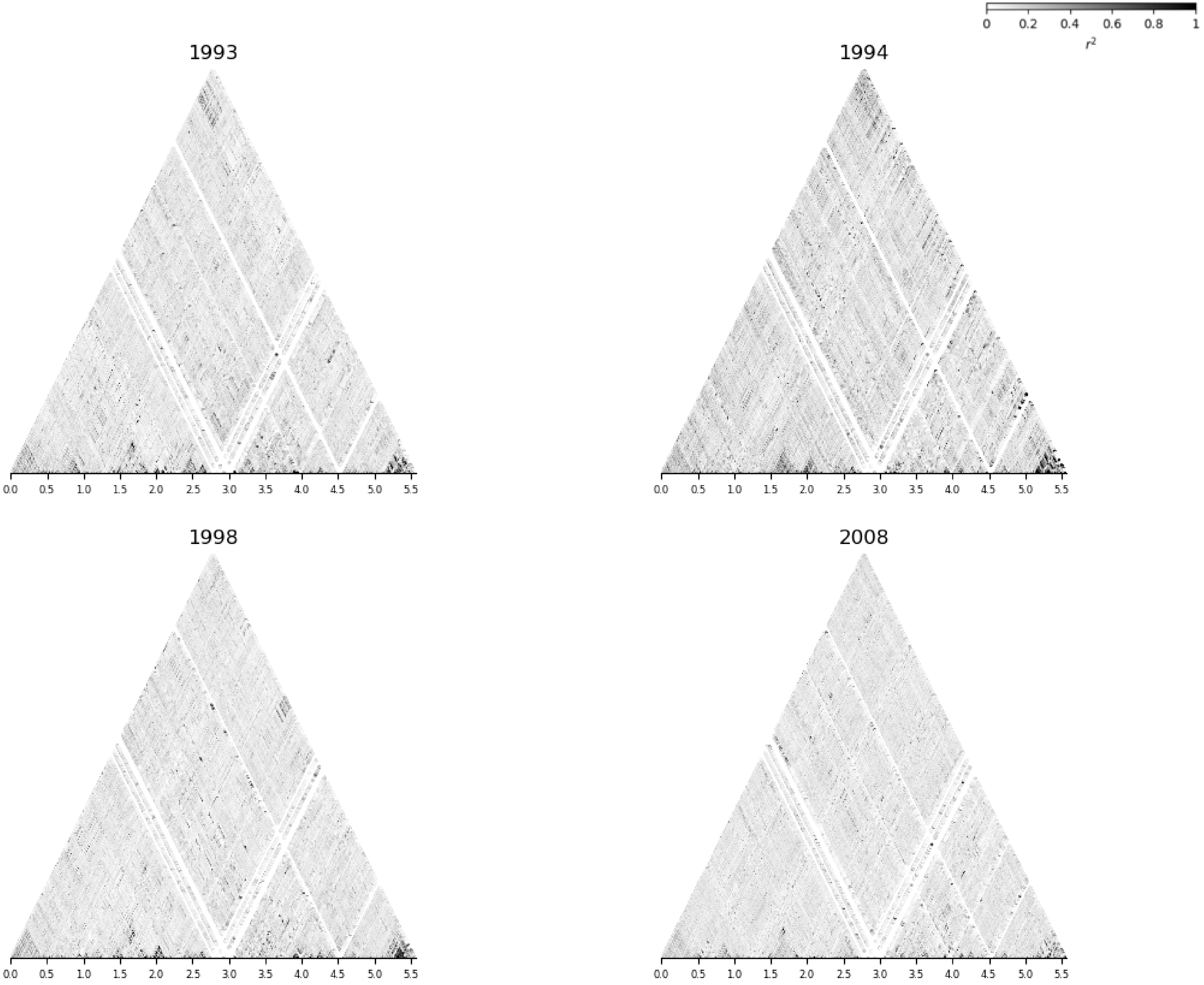
0
0.2
0.4
0.6
0.8
1
$r^2$
1993
1994
1998
2008
0.0
0.5
1.0
1.5
2.0
2.5
3.0
3.5
4.0
4.5
5.0
5.5

chr12

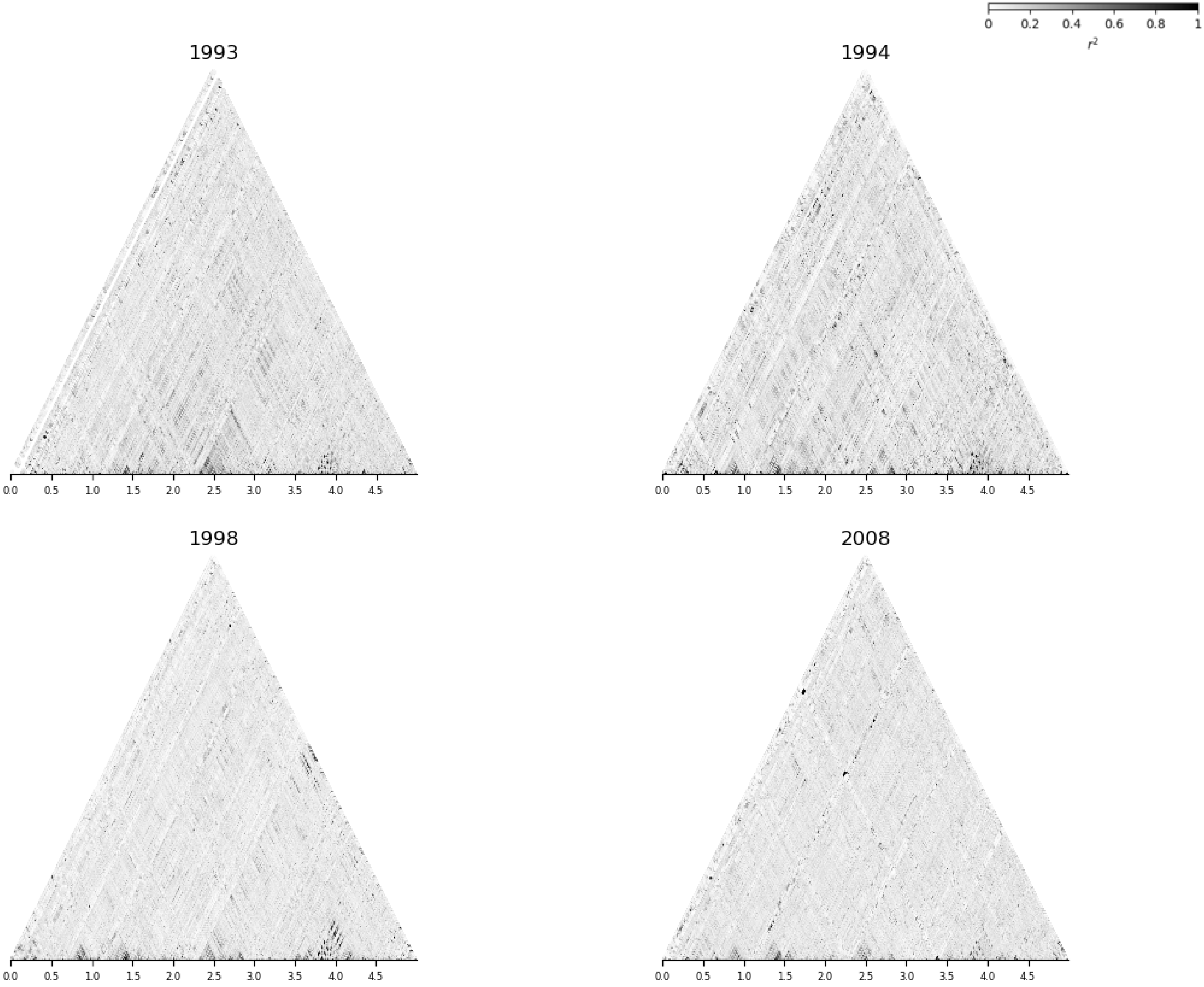

chr13

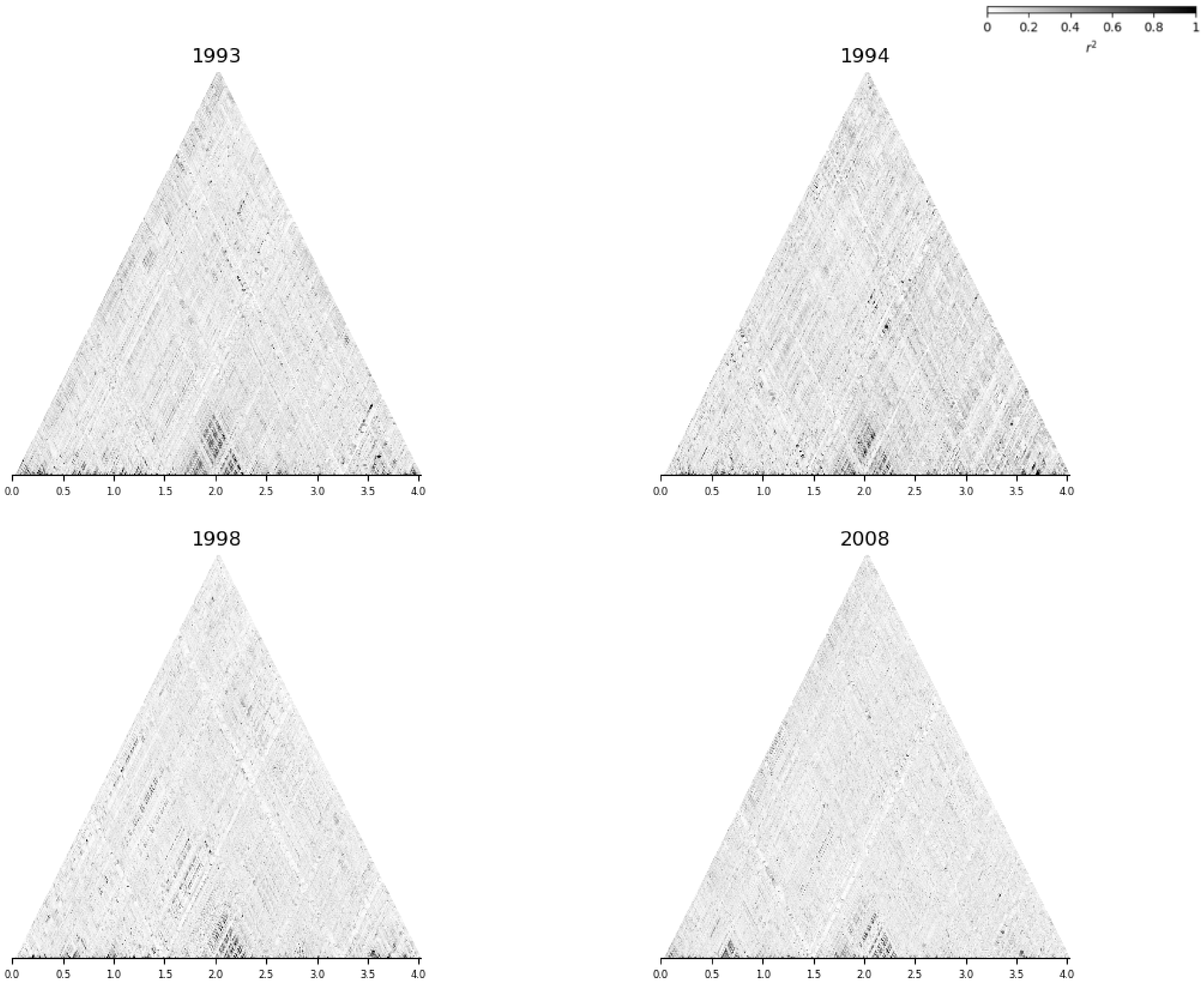

#### chr14

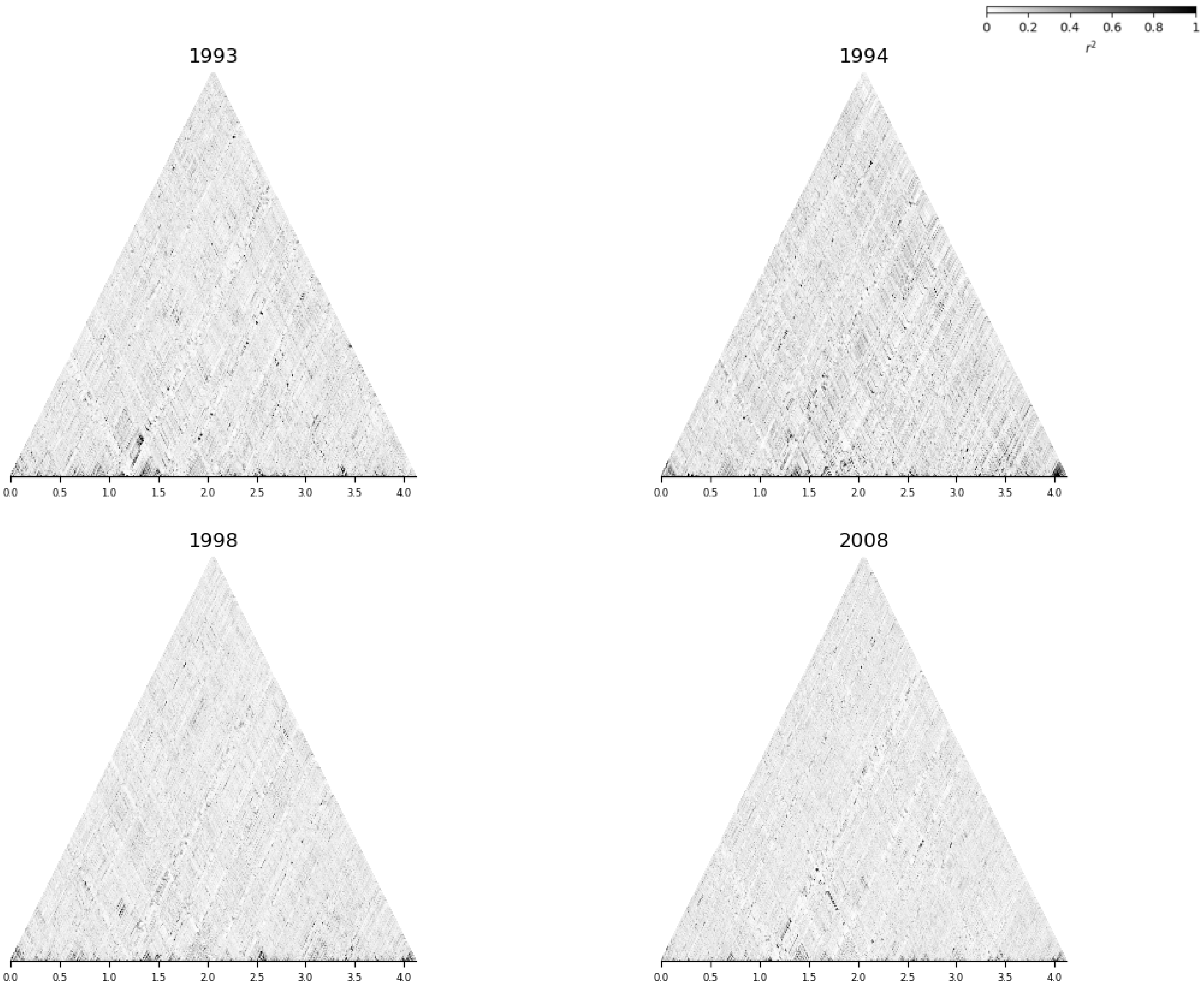

chr15

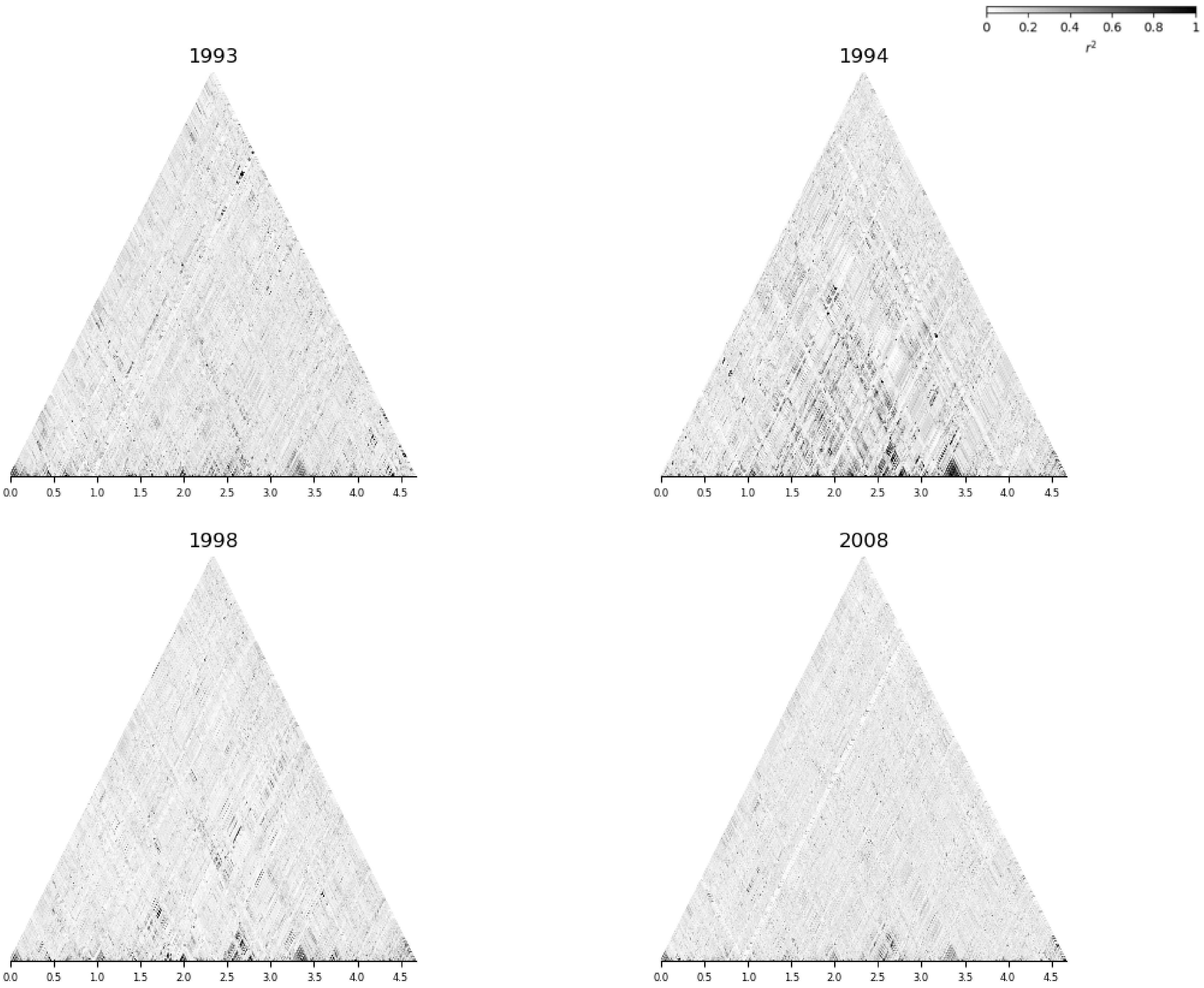

chr16

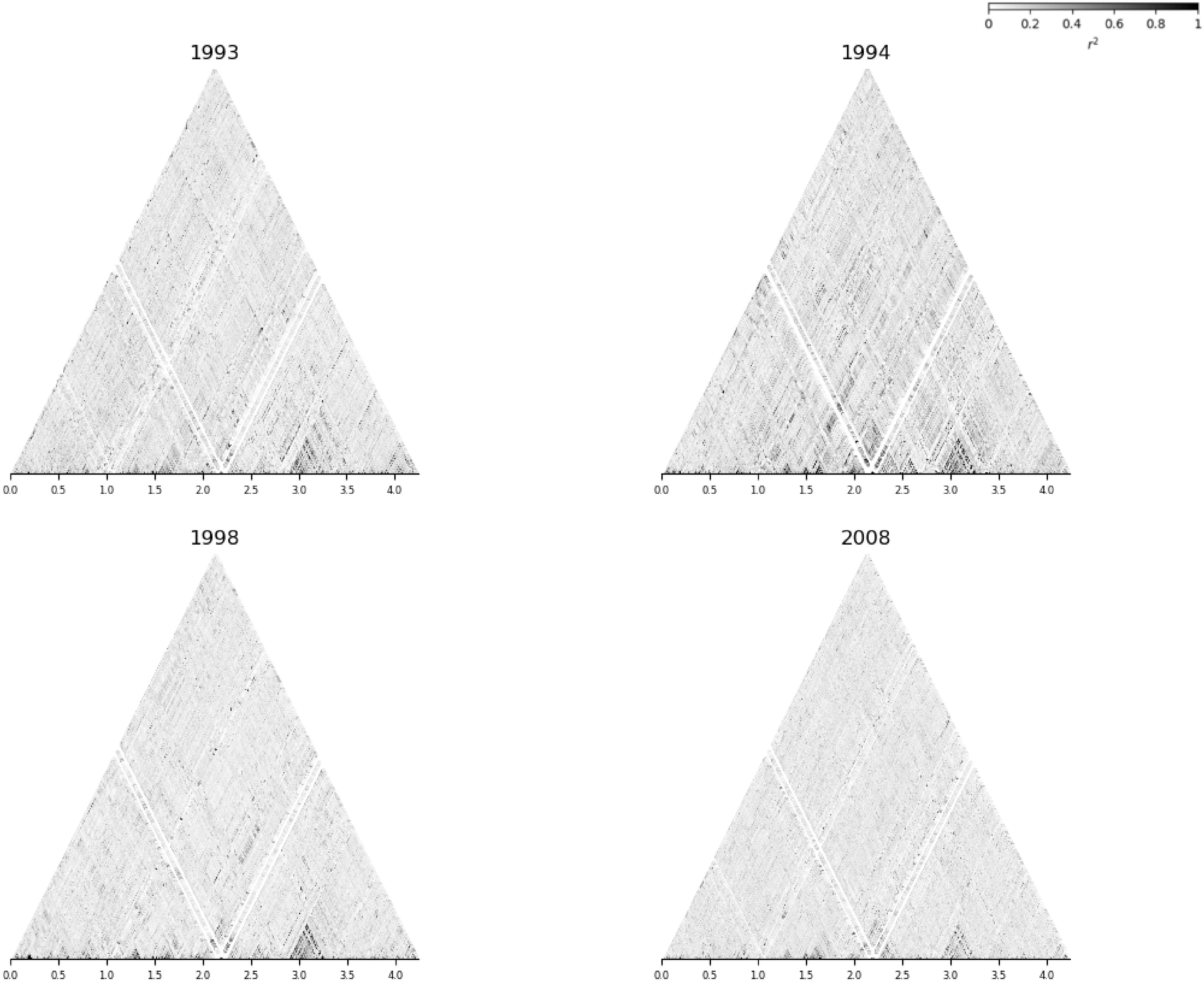

#### chr17

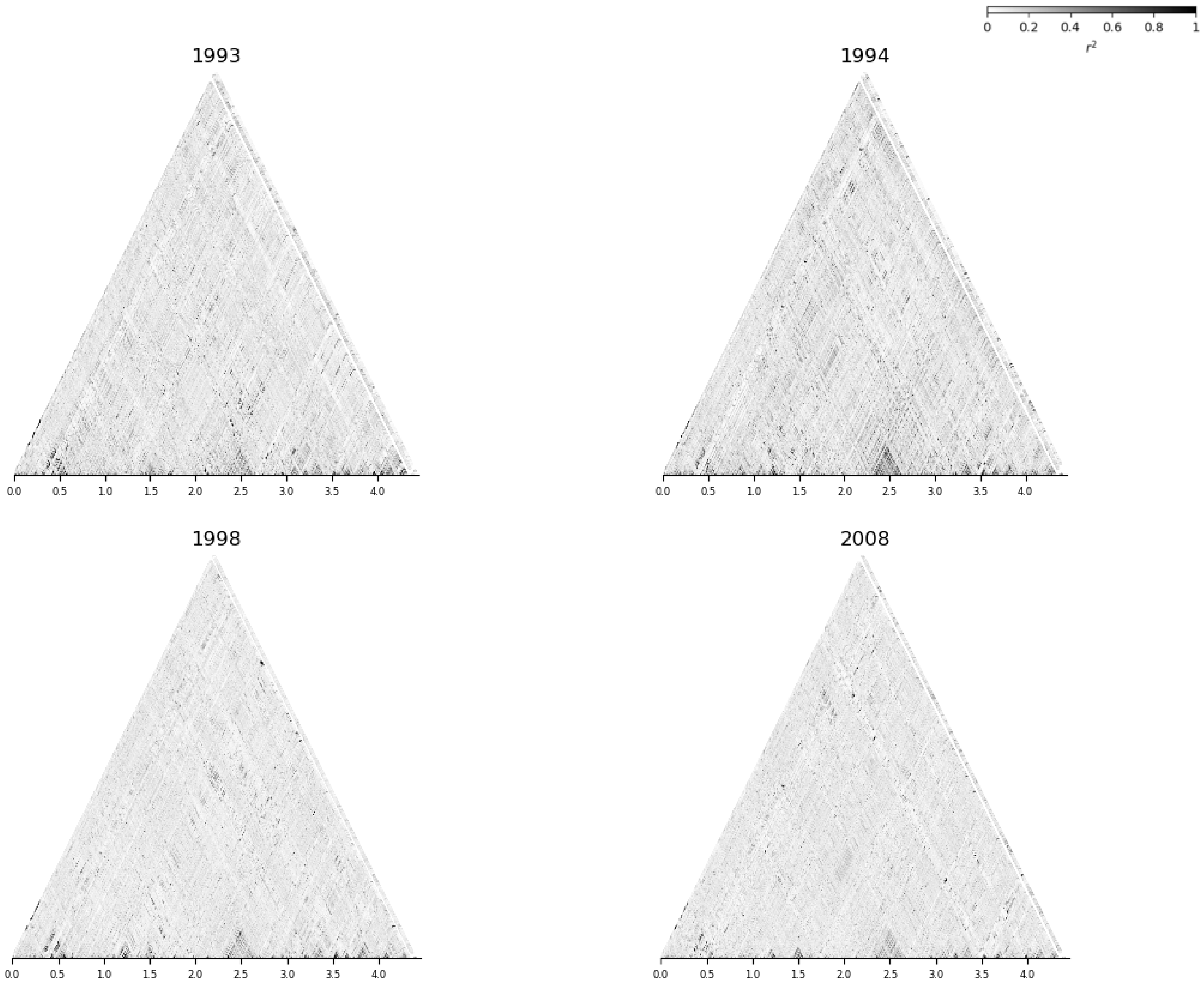

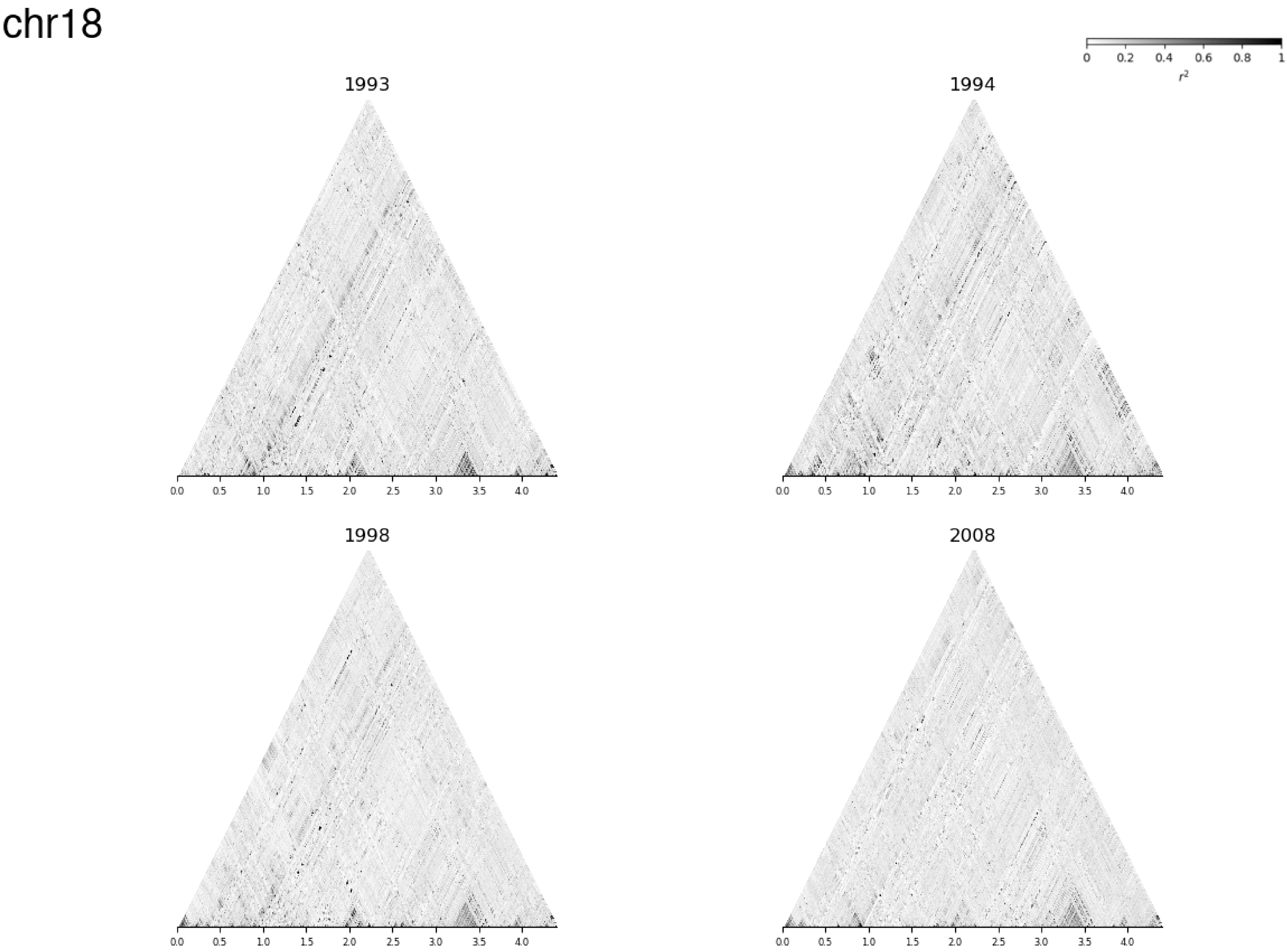
chr18
0
0.2
0.4
0.6
0.8
1
$r^2$
1993
1994
1998
2008
0.0
0.5
1.0
1.5
2.0
2.5
3.0
3.5
4.0

### Additional file 07

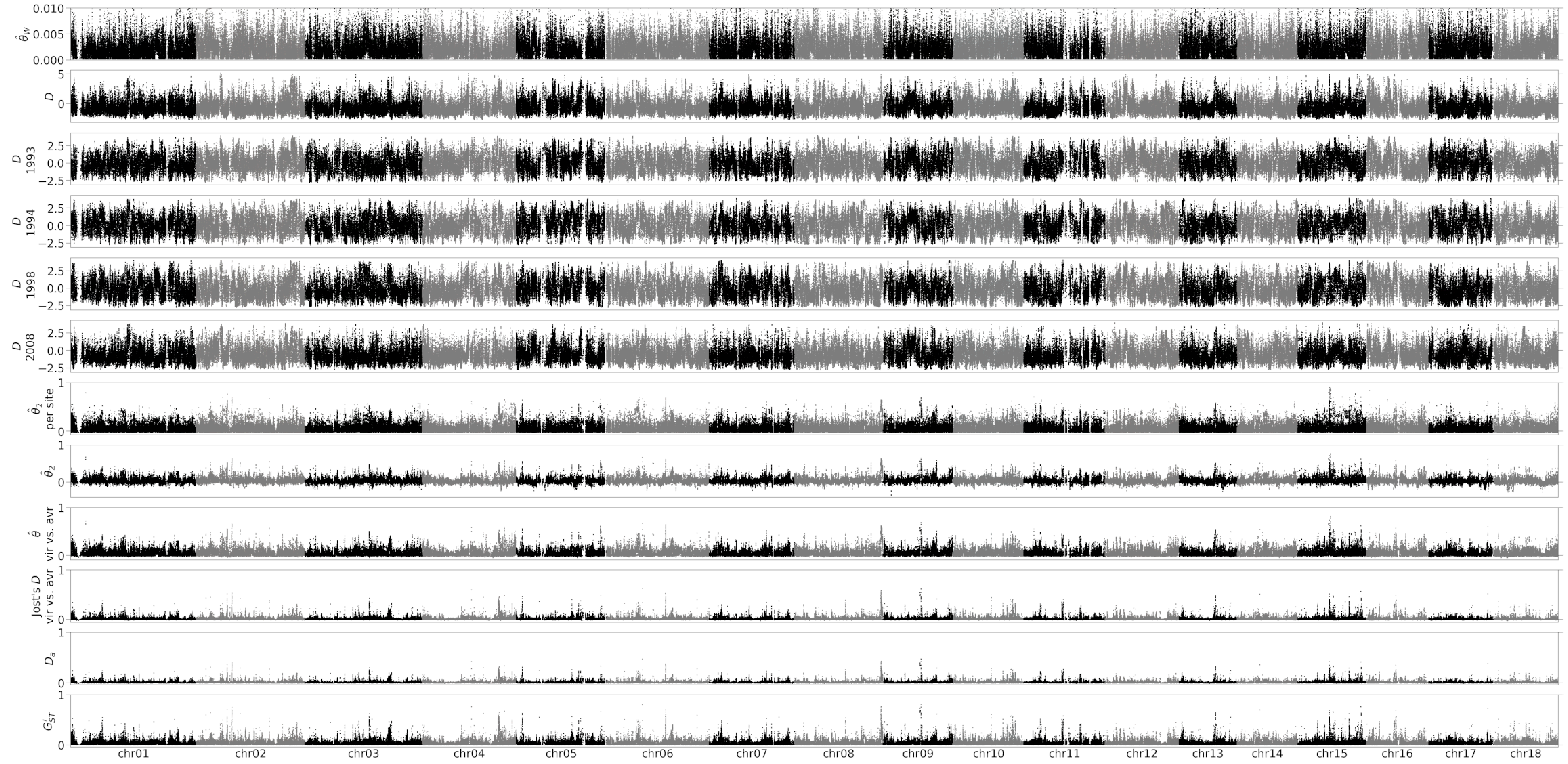

### Additional file 09

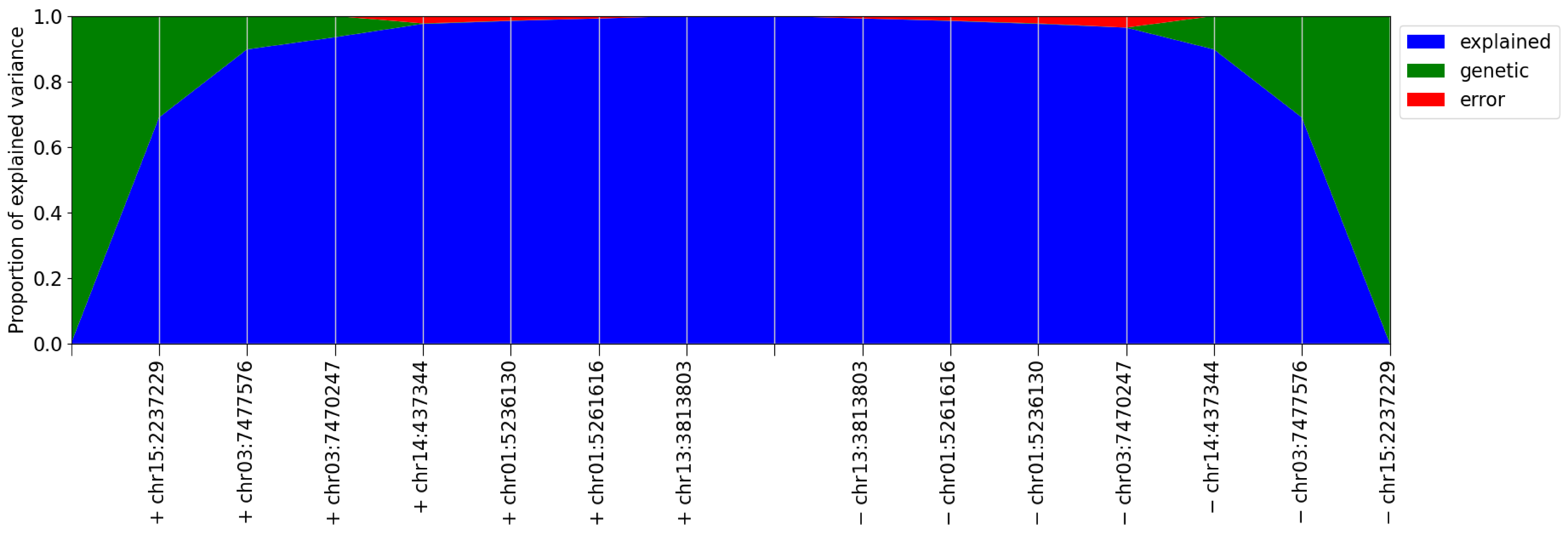

### Additional file 10

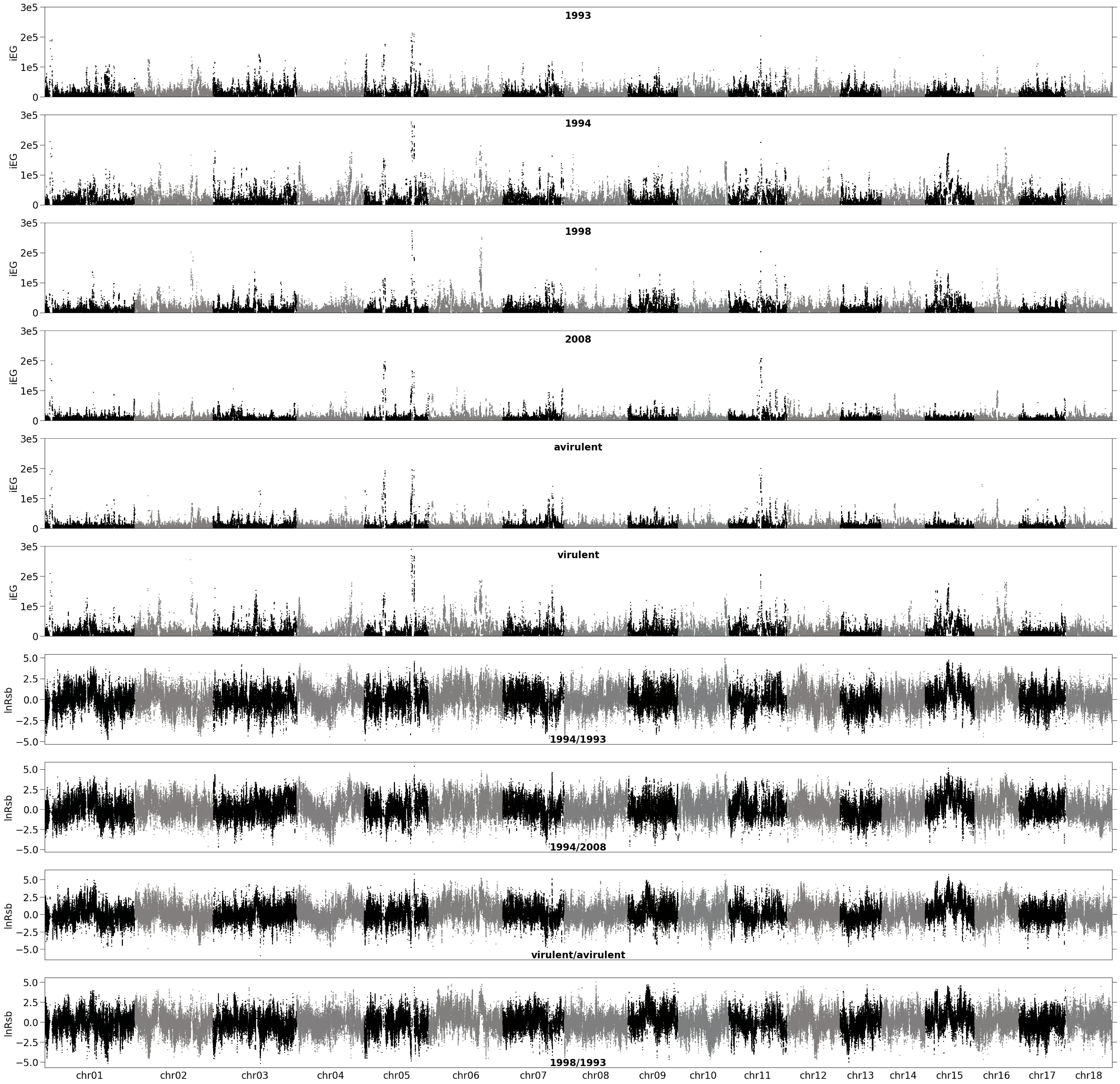

### Additional file 11

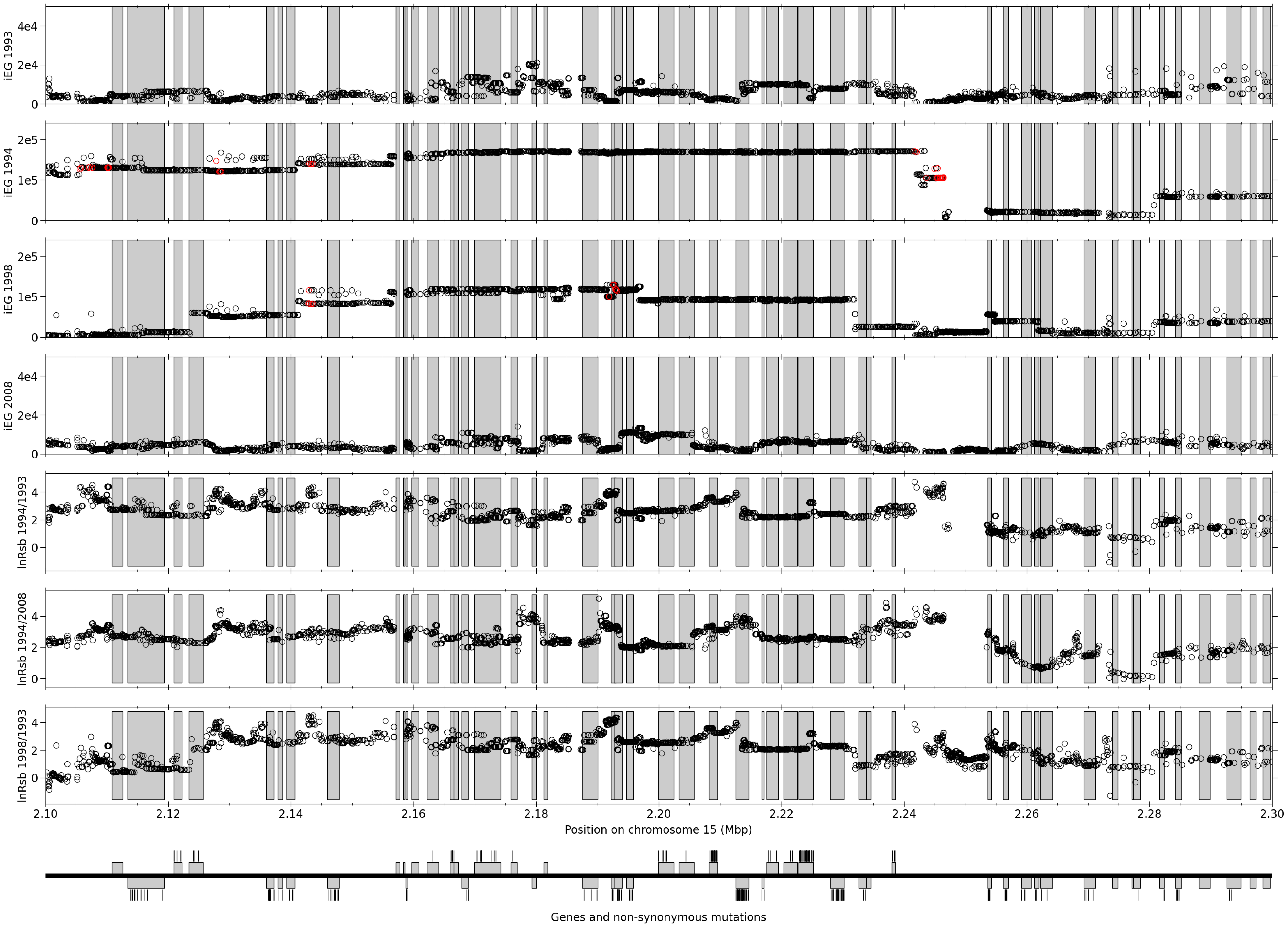
