## Additional file 05 for "Genomic signatures of a major adaptive event in the pathogenic fungus *Melampsora larici-populina*"

### Linkage disequilibrium decay

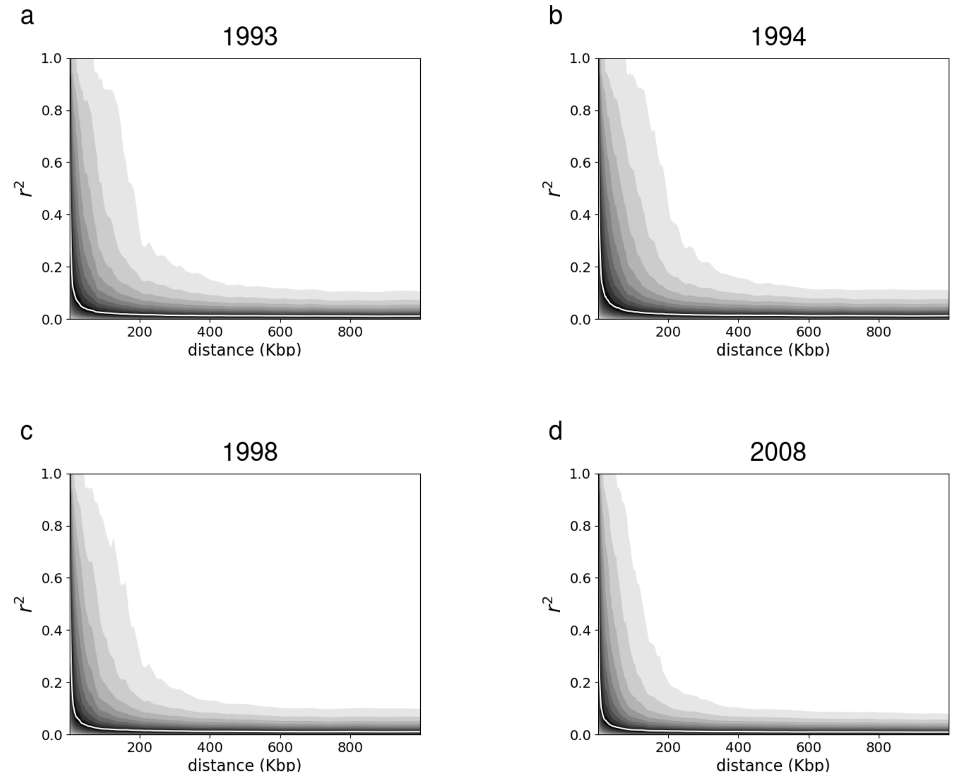


**Figure S2**. Pairwise values of $r^2$ between all pairs of site (excluding pairs of sites at a distance larger than 1Mbp and sites with more than 10% of missing data and a main allele at a frequency over 75% over the whole dataset). Pairs of sites from the 18 chromosomes are considered together. The $r^2$ values are smoothed over windows of 1000 bp with respect to the distance axis. Within each window, the median and dispersion statistics are computed. The following quantiles: 5%, 10%, 20%, 30%, 40%, 60%, 70%, 80% and 90% (shades of grey) and the median (white line) are represented.

**Table S5**. Linkage disequilibrium decay statistics. Linkage disequilibrium decay is considered as the average in 1000-bp windows based on pairwise distance. $d_t$: midpoint of the first window for which the average $r^2$ is below threshold $t$. ${r^2}_d$: average $r^2$ at window with midpoint $d$.

| Statistic | 1993 | 1994 | 1998 | 2008 |
|---|---|---|---|---|
| $d_{0.6}$ | 1500 | 1500 | 1500 | 1500 |
| $d_{0.5}$ | 2500 | 2500 | 2500 | 2500 |
| $d_{0.4}$ | 5500 | 6500 | 5500 | 4500 |
| $d_{0.3}$ | 15500 | 18500 | 13500 | 11500 |
| $d_{0.2}$ | 43500 | 59500 | 37500 | 32500 |
| $d_{0.1}$ | 145500 | 164500 | 129500 | 94500 |
| ${r^2}_{10K}$ | 0.3324 | 0.3525 | 0.3283 | 0.3093 |
| ${r^2}_{25K}$ | 0.2532 | 0.2725 | 0.2441 | 0.2275 |
| ${r^2}_{50K}$ | 0.1937 | 0.2148 | 0.1754 | 0.1667 |
| ${r^2}_{75K}$ | 0.1532 | 0.1723 | 0.1413 | 0.1251 |
| ${r^2}_{100K}$ | 0.1257 | 0.1459 | 0.1158 | 0.0955 |
| ${r^2}_{250K}$ | 0.0589 | 0.0634 | 0.0535 | 0.0418 |
| ${r^2}_{500K}$ | 0.0310 | 0.0349 | 0.0296 | 0.0240 |
