## Additional file 04 for "Genomic signatures of a major adaptive event in the pathogenic fungus *Melampsora larici-populina*"

### Summary statistics

**Table S3**. List of statistics computed in this study.

| Name | Definition |
|---|---|
| $S$ | Number of polymorphic sites |
| $\hat{\theta}_W$ | Watterson's estimator of $\theta = 4N\mu$ (expressed per exploitable site) |
| $\pi$ | Nucleotide diversity (expressed per exploitable site) |
| $D$ | Tajima's $D$ |
| $D^*$ | Fu and Li's $D$ without outgroup |
| $F^*$ | Fu and Li's $F$ without outgroup |
| $F_{\mathrm{IS}}$ | Individual fixation index |
| $D_a$ | Pairwise population net divergence |
| $G'_{\mathrm{ST}}$ | Hedrick's fixation index |
| $D_j$ | Jost's differentiation index |
| $\widehat{f}$ | Weir and Cockerham's individual fixation index estimator |
| $\widehat{\theta}$ | Weir and Cockerham's population fixation index estimator |
| $\widehat{\theta}_2$ | Weir and Cockerham's cluster fixation index estimator |
| $\widehat{F}$ | Weir and Cockerham's fixation index estimator |

**Table S4**. Genome-wide statistics. The table is broken up according to whether it is computed on the whole dataset (**a**), on individual samples (**b-e**), on the virulent (**f**) or avirulent (**g**) isolates, on the divergence between the latter two (**h**) or between all pairs of samples (**i-n**). The computed statistics are listed in **Table S3** above. Note that Weir and Cockerham's statistics can be computed while taking account the sample structure only (providing $\widehat{\theta}$) or both sample and individual structures (providing $\widehat{f}$, $\widehat{\theta}$ and $\widehat{F}$) or all of cluster, sample and individual structures (providing $\widehat{f}$, $\widehat{\theta_1}$, $\widehat{\theta_2}$ and $\widehat{F}$). The three analyses have been conducted for the whole sample and the former two have been conducted for all pairwise comparisons. **C.I. Subsampling**: 95% confidence interval based on random subsampling of sites. **Average**: average over genomic windows. **W. Average**: average weighted by the number of exploitable site per window. **C. I. Windows**: 95% confidence interval based on genomic windows.

Total number of sites: 82300004.

Variable sites: 1125506.

**a**. Whole dataset.

| **Stat** | **Value** | **C. I. Subsampling** | | **Average** | **W. Average** | **C. I. Windows** | |
|---|---|---|---|---|---|---|---|
| $\hat{\theta}_W$ | +0.002476 | +0.002472 | +0.002481 | +0.002436 | +0.002483 | +0.000432 | +0.005563 |
| $\pi$ | +0.001992 | +0.001987 | +0.001997 | +0.001982 | +0.001996 | +0.000171 | +0.006128 |
| $D$ | -0.650005 | -0.655005 | -0.645270 | -0.619118 | -0.647653 | -2.180874 | +2.276388 |
| $D^*$ | -0.340651 | -0.352694 | -0.328783 | -0.408848 | -0.405606 | -3.714620 | +2.015139 |
| $F^*$ | -0.582660 | -0.591804 | -0.573439 | -0.603446 | -0.616937 | -3.500837 | +2.136723 |
| $F_{\mathrm{IS}}$ | +0.054841 | +0.054161 | +0.055531 | +0.075684 | +0.071299 | -0.297901 | +0.436718 |
| $G'_{\mathrm{ST}}$ | +0.086054 | +0.085882 | +0.086219 | +0.087899 | +0.086458 | +0.030536 | +0.228255 |
| $D_j$ | +0.016099 | +0.016028 | +0.016162 | +0.016778 | +0.016243 | -0.000136 | +0.073460 |
| $\widehat{\theta}$ | +0.077110 | +0.076877 | +0.077344 | +0.074327 | +0.074125 | +0.000653 | +0.212405 |
| $\widehat{f}$ | -0.004733 | -0.005388 | -0.004042 | +0.020386 | +0.015734 | -0.344620 | +0.384692 |
| $\widehat{\theta}$ | +0.077119 | +0.076888 | +0.077358 | +0.073700 | +0.073638 | +0.001953 | +0.210767 |
| $\widehat{F}$ | +0.072750 | +0.072065 | +0.073455 | +0.091595 | +0.087350 | -0.289643 | +0.454807 |
| $\widehat{f}$ | -0.004733 | -0.005388 | -0.004042 | +0.020386 | +0.015734 | -0.344620 | +0.384692 |
| $\widehat{\theta_1}$ | +0.090084 | +0.089789 | +0.090385 | +0.084330 | +0.084311 | +0.002518 | +0.246933 |
| $\widehat{\theta_2}$ | +0.042136 | +0.041811 | +0.042442 | +0.036443 | +0.036416 | -0.055384 | +0.199174 |
| $\widehat{F}$ | +0.085777 | +0.085085 | +0.086503 | +0.101929 | +0.097735 | -0.281515 | +0.471382 |

(Table continues on next page.)

**Table S4** (continued).

**b**. Sample 1993.

| Stats | Value | C. I. Subsampling | | Average | W. Average | C. I. Windows | |
|---|---|---|---|---|---|---|---|
| $S$ | 669871 | 334142 | 335760 | 36.63 | 38.88 | 2 | 109 |
| $\hat{\theta}_W$ | +0.001991 | +0.001986 | +0.001995 | +0.001980 | +0.001993 | +0.000193 | +0.005587 |
| $\pi$ | +0.001928 | +0.001922 | +0.001933 | +0.001932 | +0.001932 | +0.000131 | +0.006528 |
| $D$ | -0.122079 | -0.127866 | -0.116204 | -0.110515 | -0.123895 | -2.227906 | +2.499228 |
| $D^*$ | +0.213560 | +0.205082 | +0.221417 | +0.112990 | +0.113846 | -3.939086 | +1.797715 |
| $F^*$ | +0.114426 | +0.106054 | +0.122531 | +0.045479 | +0.040543 | -3.916013 | +2.212475 |
| $F_{\mathrm{IS}}$ | -0.014783 | -0.015610 | -0.013900 | +0.004631 | +0.003206 | -0.392796 | +0.468600 |

**c**. Sample 1994.

| Stats | Value | C. I. Subsampling | | Average | W. Average | C. I. Windows | |
|---|---|---|---|---|---|---|---|
| $S$ | 607716 | 303083 | 304629 | 33.30 | 35.17 | 2 | 103 |
| $\hat{\theta}_W$ | +0.001819 | +0.001815 | +0.001824 | +0.001826 | +0.001823 | +0.000146 | +0.005359 |
| $\pi$ | +0.001844 | +0.001838 | +0.001849 | +0.001871 | +0.001848 | +0.000109 | +0.006603 |
| $D$ | +0.051762 | +0.045390 | +0.057898 | +0.025761 | +0.007593 | -2.225096 | +2.706218 |
| $D^*$ | +0.326988 | +0.318329 | +0.334850 | +0.205115 | +0.200379 | -3.990166 | +1.805325 |
| $F^*$ | +0.275049 | +0.266550 | +0.282660 | +0.172147 | +0.161211 | -3.976891 | +2.294721 |
| $F_{\mathrm{IS}}$ | +0.019227 | +0.018233 | +0.020203 | +0.039779 | +0.037420 | -0.380012 | +0.582359 |

(Table continues on next page.)

**Table S4** (continued).

**d**. Sample 1998.

| Stats | Value | C. I. Subsampling | | Average | W. Average | C. I. Windows | |
|---|---|---|---|---|---|---|---|
| $S$ | 705365 | 351869 | 353479 | 38.47 | 40.90 | 2 | 111 |
| $\hat{\theta}_W$ | +0.002040 | +0.002035 | +0.002044 | +0.002038 | +0.002051 | +0.000194 | +0.005615 |
| $\pi$ | +0.001920 | +0.001914 | +0.001925 | +0.001942 | +0.001924 | +0.000132 | +0.006636 |
| $D$ | -0.223659 | -0.229426 | -0.217799 | -0.212199 | -0.248014 | -2.338911 | +2.613768 |
| $D^*$ | -0.054533 | -0.064089 | -0.046021 | -0.123620 | -0.144969 | -4.368750 | +1.775597 |
| $F^*$ | -0.137457 | -0.146525 | -0.129039 | -0.182740 | -0.214070 | -4.307205 | +2.252848 |
| $F_{\mathrm{IS}}$ | -0.011507 | -0.012336 | -0.010601 | +0.011253 | +0.008398 | -0.357267 | +0.479383 |

**e**. Sample 2008.

| Stats | Value | C. I. Subsampling | | Average | W. Average | C. I. Windows | |
|---|---|---|---|---|---|---|---|
| $S$ | 787963 | 393123 | 394838 | 42.86 | 45.72 | 3 | 113 |
| $\hat{\theta}_W$ | +0.002309 | +0.002304 | +0.002314 | +0.002290 | +0.002318 | +0.000293 | +0.005731 |
| $\pi$ | +0.001842 | +0.001837 | +0.001847 | +0.001834 | +0.001846 | +0.000167 | +0.005803 |
| $D$ | -0.772879 | -0.777936 | -0.767943 | -0.751421 | -0.773941 | -2.374279 | +1.715386 |
| $D^*$ | -0.560061 | -0.568521 | -0.550668 | -0.597047 | -0.612679 | -4.251050 | +1.572574 |
| $F^*$ | -0.756709 | -0.765065 | -0.747745 | -0.768145 | -0.790307 | -4.239907 | +1.653139 |
| $F_{\mathrm{IS}}$ | -0.003808 | -0.004734 | -0.002931 | +0.020160 | +0.015978 | -0.358224 | +0.482529 |

(Table continues on next page.)

**Table S4** (continued).

**f**. Virulent isolates (samples 1994 and 1998 excluding 98AB07).

| **Stats** | **Value** | **C. I. Subsampling** | | **Average** | **W. Average** | **C. I. Windows** | |
|---|---|---|---|---|---|---|---|
| $S$ | 778369 | 388317 | 390003 | 42.32 | 45.13 | 2 | 119 |
| $\hat{\theta}_W$ | +0.001969 | +0.001964 | +0.001973 | +0.001956 | +0.001975 | +0.000213 | +0.005227 |
| $\pi$ | +0.001892 | +0.001887 | +0.001897 | +0.001909 | +0.001896 | +0.000130 | +0.006485 |
| $D$ | -0.137662 | -0.143473 | -0.131767 | -0.133779 | -0.165148 | -2.248739 | +2.947506 |
| $D^*$ | +0.263343 | +0.253424 | +0.272994 | +0.096189 | +0.090537 | -4.272038 | +2.007288 |
| $F^*$ | +0.117403 | +0.108670 | +0.125718 | +0.009336 | -0.009499 | -4.046092 | +2.587428 |
| $F_{\mathrm{IS}}$ | +0.017480 | +0.016692 | +0.018322 | +0.038326 | +0.035079 | -0.342161 | +0.481399 |

**g**. Avirulent isolates (samples 1993 and 2008 plus 98AB07).

| **Stats** | **Value** | **C. I. Subsampling** | | **Average** | **W. Average** | **C. I. Windows** | |
|---|---|---|---|---|---|---|---|
| $S$ | 982421 | 490248 | 492203 | 53.19 | 57.01 | 3 | 133 |
| $\hat{\theta}_W$ | +0.002463 | +0.002458 | +0.002468 | +0.002427 | +0.002469 | +0.000380 | +0.005757 |
| $\pi$ | +0.001965 | +0.001960 | +0.001970 | +0.001949 | +0.001969 | +0.000181 | +0.005986 |
| $D$ | -0.712868 | -0.717638 | -0.708139 | -0.689691 | -0.711732 | -2.247464 | +1.896603 |
| $D^*$ | -0.359659 | -0.370062 | -0.350094 | -0.435772 | -0.439481 | -4.017466 | +1.797438 |
| $F^*$ | -0.608516 | -0.617427 | -0.600311 | -0.639747 | -0.653999 | -3.881742 | +1.849436 |
| $F_{\mathrm{IS}}$ | +0.026484 | +0.025712 | +0.027234 | +0.047628 | +0.044101 | -0.322963 | +0.422993 |

(Table continues on next page.)

**Table S4** (continued).

**h**. Comparison of virulent and avirulent isolates.

| Stats | Value | C. I. Subsampling | | Average | W. Average | C. I. Windows | |
|---|---|---|---|---|---|---|---|
| $G'_{\mathrm{ST}}$ | +0.041722 | +0.041588 | +0.041853 | +0.042724 | +0.041892 | +0.008194 | +0.149701 |
| $D_a$ | +0.009884 | +0.009834 | +0.009935 | +0.010270 | +0.009940 | -0.000679 | +0.051457 |
| $D_j$ | +0.014134 | +0.014058 | +0.014212 | +0.014749 | +0.014225 | -0.001156 | +0.078791 |
| $\widehat{\theta}$ | +0.065709 | +0.065416 | +0.065999 | +0.060359 | +0.060291 | -0.004655 | +0.218294 |
| $\widehat{f}$ | +0.021769 | +0.021091 | +0.022451 | +0.045695 | +0.041216 | -0.325480 | +0.407449 |
| $\widehat{\theta}$ | +0.065409 | +0.065113 | +0.065701 | +0.059691 | +0.059691 | -0.005132 | +0.218119 |
| $\widehat{F}$ | +0.085754 | +0.085064 | +0.086478 | +0.101879 | +0.097716 | -0.281200 | +0.472106 |

**i**. Comparison of 1993 and 1994 samples.

| Stats | Value | C. I. Subsampling | | Average | W. Average | C. I. Windows | |
|---|---|---|---|---|---|---|---|
| $G'_{\mathrm{ST}}$ | +0.082106 | +0.081878 | +0.082355 | +0.084620 | +0.083679 | +0.015803 | +0.276874 |
| $D_a$ | +0.015537 | +0.015460 | +0.015621 | +0.015978 | +0.015574 | -0.002009 | +0.080373 |
| $D_j$ | +0.021667 | +0.021553 | +0.021783 | +0.022342 | +0.021756 | -0.003747 | +0.115609 |
| $\widehat{\theta}$ | +0.102349 | +0.101930 | +0.102803 | +0.094845 | +0.095295 | -0.012983 | +0.327010 |
| $\widehat{f}$ | +0.000452 | -0.000238 | +0.001200 | +0.021709 | +0.019064 | -0.372080 | +0.442447 |
| $\widehat{\theta}$ | +0.101871 | +0.101448 | +0.102334 | +0.094006 | +0.094537 | -0.013640 | +0.325646 |
| $\widehat{F}$ | +0.102277 | +0.101466 | +0.103111 | +0.112617 | +0.110812 | -0.301153 | +0.534581 |

(Table continues on next page.)

**Table S4** (continued).

**j**. Comparison of 1993 and 1998 samples.

| **Stats** | **Value** | **C. I. Subsampling** | | **Average** | **W. Average** | **C. I. Windows** | |
|---|---|---|---|---|---|---|---|
| $G'_{\mathrm{ST}}$ | +0.073304 | +0.073061 | +0.073508 | +0.075666 | +0.074576 | +0.013936 | +0.241558 |
| $D_a$ | +0.014249 | +0.014167 | +0.014320 | +0.014651 | +0.014262 | -0.001975 | +0.070797 |
| $D_j$ | +0.020110 | +0.019989 | +0.020210 | +0.020703 | +0.020140 | -0.003516 | +0.103475 |
| $\widehat{\theta}$ | +0.092789 | +0.092373 | +0.093185 | +0.085687 | +0.085883 | -0.012843 | +0.296945 |
| $\widehat{f}$ | -0.014755 | -0.015457 | -0.013969 | +0.009223 | +0.006485 | -0.352458 | +0.405155 |
| $\widehat{\theta}$ | +0.093059 | +0.092645 | +0.093449 | +0.085341 | +0.085611 | -0.012806 | +0.295458 |
| $\widehat{F}$ | +0.079677 | +0.078870 | +0.080502 | +0.092568 | +0.090371 | -0.291398 | +0.508054 |

**k**. Comparison of 1993 and 2008 samples.

| **Stats** | **Value** | **C. I. Subsampling** | | **Average** | **W. Average** | **C. I. Windows** | |
|---|---|---|---|---|---|---|---|
| $G'_{\mathrm{ST}}$ | +0.059423 | +0.059259 | +0.059593 | +0.060299 | +0.059573 | +0.013816 | +0.193645 |
| $D_a$ | +0.010859 | +0.010805 | +0.010919 | +0.011226 | +0.010981 | -0.002113 | +0.058150 |
| $D_j$ | +0.015213 | +0.015130 | +0.015299 | +0.015770 | +0.015423 | -0.003739 | +0.086470 |
| $\widehat{\theta}$ | +0.073794 | +0.073466 | +0.074142 | +0.067696 | +0.067889 | -0.013772 | +0.252472 |
| $\widehat{f}$ | -0.010467 | -0.011235 | -0.009745 | +0.013274 | +0.009654 | -0.362864 | +0.405640 |
| $\widehat{\theta}$ | +0.073813 | +0.073487 | +0.074168 | +0.067191 | +0.067497 | -0.013845 | +0.251165 |
| $\widehat{F}$ | +0.064118 | +0.063286 | +0.064915 | +0.079245 | +0.076280 | -0.308035 | +0.473055 |

(Table continues on next page.)

**Table S4** (continued).

**l**. Comparison of 1994 and 1998 samples.

| Stats | Value | C. I. Subsampling | | Average | W. Average | C. I. Windows | |
|---|---|---|---|---|---|---|---|
| $G'_{\mathrm{ST}}$ | +0.038632 | +0.038499 | +0.038743 | +0.040236 | +0.039228 | +0.007310 | +0.137936 |
| $D_a$ | +0.003537 | +0.003504 | +0.003566 | +0.003808 | +0.003592 | -0.004910 | +0.028909 |
| $D_j$ | +0.005186 | +0.005132 | +0.005232 | +0.005646 | +0.005294 | -0.008730 | +0.046925 |
| $\widehat{\theta}$ | +0.025026 | +0.024812 | +0.025219 | +0.024817 | +0.024517 | -0.023173 | +0.143818 |
| $\widehat{f}$ | +0.002664 | +0.001884 | +0.003515 | +0.024496 | +0.021087 | -0.355325 | +0.468478 |
| $\widehat{\theta}$ | +0.024892 | +0.024680 | +0.025088 | +0.023969 | +0.023804 | -0.026036 | +0.141530 |
| $\widehat{F}$ | +0.027489 | +0.026662 | +0.028315 | +0.047437 | +0.044051 | -0.340994 | +0.487428 |

**m**. Comparison of 1994 and 2008 samples.

| Stats | Value | C. I. Subsampling | | Average | W. Average | C. I. Windows | |
|---|---|---|---|---|---|---|---|
| $G'_{\mathrm{ST}}$ | +0.065984 | +0.065793 | +0.066190 | +0.067562 | +0.066028 | +0.014390 | +0.242362 |
| $D_a$ | +0.012844 | +0.012776 | +0.012920 | +0.013661 | +0.013026 | -0.002185 | +0.078536 |
| $D_j$ | +0.018054 | +0.017950 | +0.018166 | +0.019259 | +0.018338 | -0.003974 | +0.116701 |
| $\widehat{\theta}$ | +0.087504 | +0.087102 | +0.087948 | +0.078800 | +0.078318 | -0.013960 | +0.303681 |
| $\widehat{f}$ | +0.006556 | +0.005763 | +0.007311 | +0.029500 | +0.025203 | -0.353129 | +0.461062 |
| $\widehat{\theta}$ | +0.087255 | +0.086851 | +0.087703 | +0.077877 | +0.077522 | -0.015504 | +0.302945 |
| $\widehat{F}$ | +0.093239 | +0.092392 | +0.094091 | +0.104745 | +0.100589 | -0.289133 | +0.534864 |

(Table continues on next page.)

**Table S4** (continued).

**n**. Comparison of 1998 and 2008 samples.

| Stats | Value | C. I. Subsampling | | Average | W. Average | C. I. Windows | |
|---|---|---|---|---|---|---|---|
| $G'_{\mathrm{ST}}$ | +0.059709 | +0.059512 | +0.059901 | +0.061319 | +0.059729 | +0.012866 | +0.211005 |
| $D_a$ | +0.011901 | +0.011832 | +0.011970 | +0.012590 | +0.012012 | -0.002130 | +0.068468 |
| $D_j$ | +0.016929 | +0.016828 | +0.017031 | +0.017919 | +0.017090 | -0.003703 | +0.101293 |
| $\widehat{\theta}$ | +0.079666 | +0.079271 | +0.080052 | +0.070799 | +0.070153 | -0.013463 | +0.271782 |
| $\widehat{f}$ | -0.009007 | -0.009750 | -0.008258 | +0.016496 | +0.011882 | -0.352656 | +0.426188 |
| $\widehat{\theta}$ | +0.080244 | +0.079852 | +0.080637 | +0.070425 | +0.069898 | -0.013917 | +0.271593 |
| $\widehat{F}$ | +0.071960 | +0.071162 | +0.072810 | +0.084967 | +0.080369 | -0.293194 | +0.505265 |
